# Palaeogenomics-informed inferences of European dog admixture enables scalable dingo conservation

**DOI:** 10.64898/2026.04.08.717357

**Authors:** Shyamsundar Ravishankar, Nhi Chau Nguyen, Leonard Taufik, Nathan M. Michielsen, Anders Bergström, Raymond Tobler, Damien Fordham, Anna Brüniche-Olsen, Carsten Rahbek, Bastien Llamas, Yassine Souilmi

## Abstract

Dingoes, mainland Australia’s sole terrestrial apex mammal for over 3,000 years, are important components of many ecosystems and Indigenous cultural heritage. Yet conflicts with farmers over livestock predation following European colonisation led to widespread lethal control. These measures are further reinforced by perceptions of hybrid ancestry with European dogs. Accurate estimation of European dog ancestry is therefore essential for effective conservation, but existing tests yield highly conflicting results.

Leveraging pre-colonial dingo palaeogenomes and a robust ancestry modelling framework, we reassess the genetic ancestry of contemporary populations. Our approach corrects limitations and biases in existing methods, producing consistent estimates even with as few as 10,000 genome-wide transversion genetic markers. Accounting for admixture uncovers population structure that has persisted for over two millennia and reveals patterns of genetic admixture coinciding with human activity during the colonial era.

This study underscores the value of palaeogenomes as a vital conservation tool, offering insights unattainable from modern DNA alone. By clarifying ancestry and population structure, our study offers a robust foundation for effective regionally informed dingo management across Australia.

## Introduction

Dingoes fulfil a unique ecological role in Australian ecosystems (Newsome et al. 2015; Wooster et al. 2024), becoming the sole terrestrial apex predator on the mainland (and several offshore islands) shortly after their arrival over 3,000 years before present (BP) (Corbett, L. K. 1985). Culturally, they hold great significance to many Indigenous Australians, often appearing in ancestral songlines as key contributors to ecological and cultural balance (Koungoulos et al. 2023). However, dingoes have been involved in ongoing conflicts with livestock farmers since the early colonial period (1800s), resulting in the implementation of dingo management measures across Australia. Widely employed management measures include culling and territorial restraint, the latter most notably attested in the erection of a 5,615 km fence across Australia to restrict dingo access to farming areas (**Fig. 1b**). Complicating this management is the introduction of European dogs since the early colonial period, which led to interbreeding with dingoes in some regions. Today, free-living canines are killed and excluded under the classification of “wild dogs”—encompassing dingoes, dingo and dog hybrids, and feral dogs— to reduce livestock predation. This has prompted calls for the use of genetic tests to estimate dingo ancestry in free-living canines to prevent loss of dingo genetic diversity, especially given that both feral dogs and first generation (F1) dingo-dog hybrids are rare (Cairns et al. 2021).

**Figure 1.**
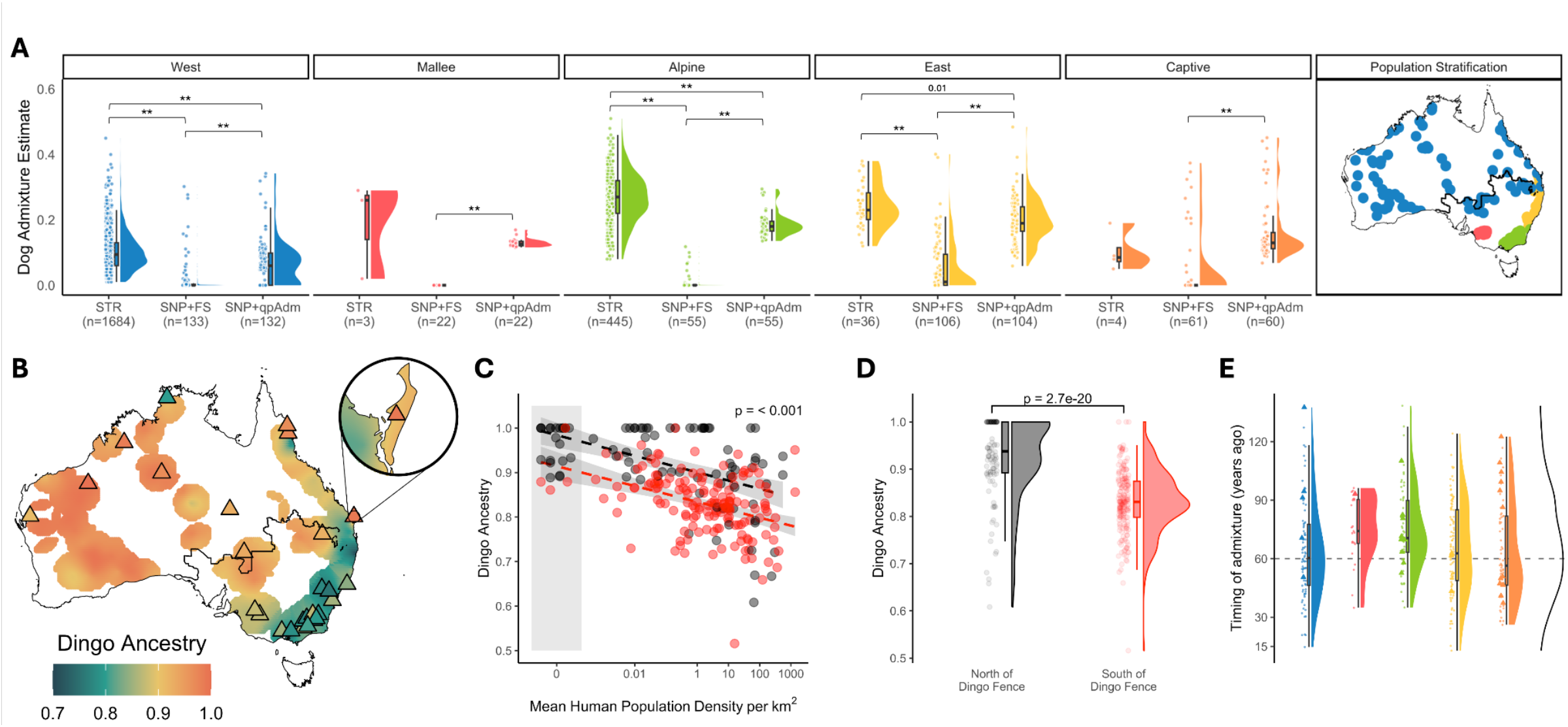
Dog Admixture Proportions. **A)** Dog admixture estimates for free-roaming and captive dingoes from around Australia using microsatellite data (STR - previously estimated in (Stephens et al. 2015)); SNP-Array data with dog admixture estimated using FastStructure program (SNP+FS - previously estimated in (Cairns et al. 2023)); and using SNP-Array data with dog admixture modelled using qpAdm as described in the ‘Methods’ (SNP+qpAdm). The populations are stratified as previously defined in (Cairns et al. 2023). ** Shows significant difference between tests (*p* < 0.01). Numbers below each label shows the number of samples used in comparison. Panels **B, C & D** use dog admixture or dingo ancestry (i.e., 1 - proportion of dog admixture) from SNP-Array and Whole Genome Sequencing (WGS) data estimated using qpAdm. **B)** Interpolated dingo ancestry across Australia. The Dingo Fence is highlighted on the map (bold black line). WGS samples are represented using triangles. For visualisation, ancestry was averaged such that there were at least two samples within 200km of each other. See **SI Section 4** for more details. **C)** Dingo admixture as a function of mean human population density within a 400 km^2^ radius. Points are colored by location relative to the Dingo Fence: north (black) and south (red). The panel is annotated with the *p*-value (< 0.001) from the beta-regression model accounting for human population density, position relative to fence and dingo ancestry. **D)** Distribution of dingo ancestry north (black) and south (red) of the Dingo Fence, showing a highly significant difference (*p* = 2.7e-20). **E)** Timing of admixture between dingoes and European dogs by population, inferred from MOSAIC (Salter-Townshend & Myers 2019) analysis of imputed SNP-Array and WGS individuals (dots and triangles, respectively) with less than 95% dingo ancestry, as estimated using qpAdm. Generation times from MOSAIC were converted to years by assuming a 3-year generation time (Scarsbrook et al. 2025). The dashed line marks 60 years ago (i.e. 1960s, assuming all samples were collected today).

The emergence of dingo ancestry as a component of management and conservation efforts across Australia has prompted the development of genomic assays that aim to accurately measure European dog ancestry proportions. However, the two most commonly used assays for inferring European dog ancestry proportions have produced dramatically different estimates, with microsatellite (STR; 24 loci) tests typically identifying extensive admixture among assayed dingoes (Wilton et al. 1999; Wilton 2001; Stephens et al. 2015), whereas the recently introduced SNP-array (195,474 SNPs) and DArTseq (2,466 SNPs) tests showed limited and almost no admixture, respectively, in the majority of free-roaming dingoes (Cairns et al. 2023; Weeks et al. 2025). Whether these discrepancies reflect differences in genetic marker ascertainment or estimation methods remains unclear. A key limitation shared by these methods is their reliance on contemporary dingo populations assumed to be free of admixture as an appropriate reference. Recent studies addressed this limitation by using pre-colonial dingo genomes, providing a more robust baseline for ancestral dingo diversity in the absence of European dog admixture (Souilmi et al. 2024; Scarsbrook et al. 2025). These studies reveal low levels (< 20%) of European dog admixture in contemporary dingoes and suggest regional variation in admixture across Australia but the relatively small number of individuals limits the generality of their findings.

Because lethal control remains a fundamental measure in dingo management across Australia, accurate quantification of European dog admixture proportions is essential to avoid detrimental consequences for dingoes, including local extinctions and inadvertent loss of ancestral genetic diversity and adaptive variation (Claridge et al. 2014). Introgressed variation from European dogs may partially alleviate inbreeding depression and contribute adaptive variation in southeastern dingoes (Scarsbrook et al. 2025), complicating efforts to manage dingoes solely on the basis of ancestry. Nevertheless, the development of appropriate conservation strategies across Australia requires a detailed and accurate understanding of dingo population structure and connectivity at regional levels, which critically requires the use of methods that are robust to different data types and not prone to bias.

Accordingly, in this study, we employ a genomic approach capable of producing robust ancestry estimates (qpAdm (Harney et al. 2021)) to precisely assess the European dog admixture levels of more than three hundred dingoes previously assayed with SNP-array or whole-genome sequencing (WGS) data across the species range. Crucially, by using recently published pre-colonial dingo palaeogenomes to characterise baseline dingo ancestry, we show that the qpAdm approach provides robust estimates with as few as 10,000 genome-wide single nucleotide polymorphisms (SNP), while also identifying the limitations in prior tests. By applying previously described local-ancestry inference methods (Scarsbrook et al. 2025) to hundreds of free-living dingoes, we additionally provide a high resolution map of modern dingo genetic diversity, revealing biogeographic population structure that are influenced by natural barriers, and anthropogenic activity following European colonisation of Australia. Taken together, our findings provide the most comprehensive study of dingo population genetic structure yet reported and reinforce the importance of using robust ancestry estimation methods for informing effective and evidence-based strategies for meaningful dingo conservation and management.

## Results

### Revised Dingo Admixture Estimates

To obtain comprehensive estimates of European dog ancestry levels in dingoes across Australia, we used the qpAdm method (Haak et al. 2015; Harney et al. 2021) to model dingo ancestry for 347 individuals that were either assayed using SNP-Array data (n = 299, genotyped at >190 thousand loci using the Axiom Canine HD Genotyping array) (Cairns et al. 2023) or had WGS (n = 48, including two dingo reference genomes) (Zhang et al. 2020; Field et al. 2022; Ballard et al. 2023; Souilmi et al. 2024; Scarsbrook et al. 2025) (**Table S2**; see **Methods and SI Section 5**). Importantly, the qpAdm approach has the advantage of being able to test specific historical admixture models with appropriate ancestry proxies—i.e. pre-colonial dingo palaeogenomes as an unadmixed baseline (Souilmi et al. 2024; Scarsbrook et al. 2025) and four high coverage German Shepherd genomes to represent European dog ancestry (Plassais et al. 2019)—while remaining robust to potential confounding factors like post-admixture drift and marker ascertainment (Harney et al. 2021; Williams et al. 2024).

We show that this ancestry modelling framework is generally robust to the choice of different European dog breeds used as sources, different ascertainment schemes (including SNP-Array data and whole-genome enrichment)—and a reduced number of loci (remaining informative at 10,000 pseudohaploid SNPs) (see **SI Section 5**). Nevertheless, we advise caution when applying qpAdm in cases where genetic information in the source and the closest target populations have been ascertained using different non-random schemes (e.g. SNP-Array and whole-genome enrichment) (see **SI Section 5.2**).

Comparing our revised qpAdm-based estimates of European dog admixture to previously reported estimates from STR (2,168 dingoes, 24 STR loci) (Stephens et al. 2015) and the 299 SNP-array assayed dingoes reveals consistent differences between the three approaches. Specifically, we observed that reported European dog ancestry levels were significantly underestimated for the SNP-array assayed dingoes (*t*-test *p-*value < 0.01) whereas the estimates from STR loci tended to significantly overestimate European dog admixture for all subgroups other than “Mallee” and “Captive” dingoes (**Fig. 1A**). Though it should be noted that comparisons between STR and SNP-Array are based on different sets of individuals. We confirm that the significantly lower European dog admixture estimates in SNP-Array assayed dingoes are not attributable to marker ascertainment, and that the SNP-Array loci are sufficiently informative to robustly distinguish between dingo and dog ancestries (**Fig. S5A, SI Section 5.4**).

To assess the impact of the inference method on ancestry estimation, we compared qpAdm estimates to ADMIXTURE, a widely used model-based clustering approach that decomposes ancestry components, being analogous to the FastStructure method used previously (Cairns et al. 2023). Similar to the FastStructure results, ADMIXTURE consistently overestimates dingo ancestry for the SNP-Array data, with similar overestimates also observed for WGS individuals. Specifically, ADMIXTURE estimates of European dog ancestry in contemporary dingoes decrease as the assumed number of ancestral components increase (**Fig. S10B**), especially in the “East” and “Alpine” subgroups. This suggests that clustering-based methods may be confounded by historical gene flow from a European dog lineage that is now ubiquitous within tested populations (Scarsbrook et al. 2025) (see **Timing of gene flow from European dogs** and **Fig. 1E**), leading to inflated estimates of dingo ancestry proportions. These findings underscore the limitations of clustering-based methods when directly estimating admixed ancestry compositions.

Taken together, our findings indicate that both genetic ascertainment and method choice can introduce limitations that exacerbate biases in estimates of European dog ancestry in dingoes. While the most robust inferences are obtained using qpAdm applied to whole-genome data, qpAdm estimates remain robust across different data types—even with reduced marker sets (**SI Section 5**). Estimates from a recently used local-ancestry inference method, MOSAIC, were consistent with qpAdm for whole-genome data but showed some discrepancies for SNP-Array data (see **Fig S8, SI Section 6.2**). By contrast, clustering-based approaches such as ADMIXTURE can produce inconsistent estimates, and we therefore caution against its use for quantifying admixture in dingoes.

### Human impact on dingo ancestry

When scrutinising the spatial distribution of our revised dingo ancestry estimates, we observed distinctive geographical patterns that suggest proximity to population centres, differences in land-use regimes, and the dingo fence most likely influences the degree of admixture with European dogs (**Fig. 1B, 1C, 1D**). To formally assess these relationships, we modelled our revised dingo ancestry estimates as a function of human population density and position relative to the dingo fence using a general linear modelling approach that accounts for spatial autocovariance, using a combination of SNP-Array and WGS sequenced dingoes (see **SI Section 2.4**). Our results show that dingo ancestry decreases significantly with increasing present-day human population density (**Fig. 1C**). We also observe a significant difference in admixture levels north and south of the fence with higher European dog admixture observed in dingoes south of the fence (**Fig. 1D**). However, it remains unclear whether this reflects a direct effect of the fence itself or is instead confounded by underlying population structure and correlated environmental variables, such as differences in lethal management programs. Long-term monitoring of dingo ancestry in Western Australia has failed to produce detectable genetic changes in admixture levels within a fenced region (Stephens et al. 2023), although the study did identify the fence as a barrier to population connectivity. Together, our results suggest that the regional variation in dingo ancestry is primarily correlated with human population density (**Table 1**).

**Table 1.** Results of the beta regression model predicting dingo ancestry as a function of log-transformed human population density, position relative to the dingo fence and a spatial random effect between regions north and south of the fence. Significant *p*-values (< 0.05) are indicated in italics.

| Variable | Estimate | Std. Error | *p*-value |
| --- | --- | --- | --- |
| Intercept | 3.79682 | 0.14494 | *< 0.001* |
| log(human density) | -0.20988 | 0.03546 | *< 0.001* |
| South of the dingo fence | -1.89804 | 0.14910 | *< 0.001* |
| MEM (spatial autocorrelation) | -0.18410 | 0.06071 | *0.002* |
| Spatial random effect | $3.181 \times 10^{-10}$ | $1.784 \times 10^{-5}$ | - |

### Timing of gene flow from European dogs

In agreement with recent inferences from WGS dingoes (Scarsbrook et al. 2025), we find that admixture from European dogs occurred predominantly during the 1950s and 1960s (approximately 20–25 generations ago, assuming an average 3-year generation time) (**Fig. 1E**). Interestingly, despite our ancestry estimates indicating regional differences in admixture levels (**Fig. 1B**), the timing of gene flow from European dogs appears to have been broadly ubiquitous across the continent (**Fig. 1E, S9A, S9B**). This suggests that dingoes across Australia experienced varying rates of admixture from European dogs, with southeastern dingoes likely experiencing higher rates than the rest. We also observe individuals in western Australia showing more recent European dog gene flow (<6 generations ago) while retaining low overall European dog ancestry (<10%). These cases indicate that even when admixture is recent, subsequent generations can largely restore dingo ancestry, provided unadmixed mates remain available (**Fig. S9C**).

### Population structure

A recent study of dingo genetic history showed that contemporary dingo population structure is broadly divided across two distinct subgroups situated on either side of the Great Dividing Range (Souilmi et al. 2024), with those to the west having a stronger affinity to ancient dingoes from Nullarbor, and the eastern group showing more affinity to the ancient Curracurrang dingoes. To see if these results apply more generally throughout Australia, we evaluated the ancestral affinities of 285 SNP-Array and WGS dingoes after masking European dog ancestry, using *f*_4_ statistics to estimate the shared genetic drift (see **SI Section 3.3**). Our results confirm that this population subdivision holds broadly across modern dingoes distributed throughout Australia, with the transition between the western and eastern groups coinciding broadly with the western hinterlands of the Great Dividing Range and the Murray-Darling river system, although denser sampling across these regions will be required to resolve the precise role of these features (**Fig. 2A**).

**Figure 2.**
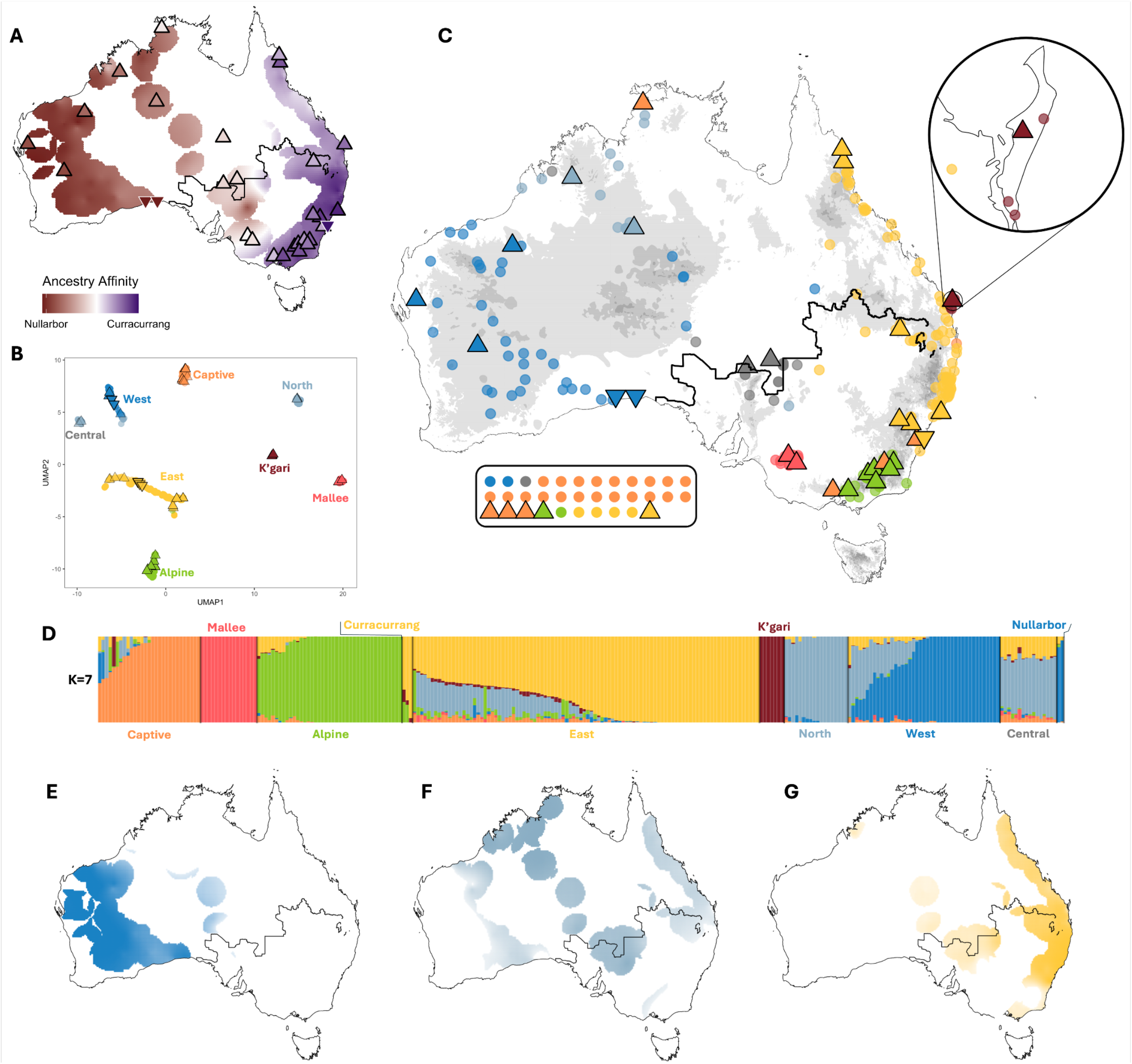
Ancestral Dingo Population Structure. All panels use genomic data from WGS and SNP-Array dingoes. **A)** Normalised outgroup *f*_4_-values of the form *f*_4_*(X, CoyoteCalifornia; Y, AndeanFox)*, where *X* is the test modern dingo individuals with dog ancestry masked (See **SI Methods**) and *Y* represents either the ‘Nullarbor or Curracurrang lineages. Maroon indicates a stronger affinity towards the Nullarbor lineage and purple the Curracurrang lineage. Triangles and inverted triangles in all panels represent modern and ancient WGS dingoes, respectively. Dots in panel **B** are SNP-Array samples. **B)** UMAP plot based on the top 10 principal components (PCs), calculated using only ancestral dingo segments from 283 unrelated individuals. Samples are grouped based on separation in UMAP space and labelled by their geographic provenance. Samples are colored by the most prominent ancestry inferred by ADMIXTURE at K=7, with transparency scaled according to the weight of that ancestral component. **C)** Geographical distribution of samples coloured by population labels from panel **B**. Sample shapes follow the same pattern as panel **B**. Samples within the box are samples that do not have coordinates and mostly represent individuals in captivity. **D)** ADMIXTURE results for the model with the lowest cross-validation error (K=7, see **Fig. S15**). Samples are grouped and labelled as in panel **B**, with the exception of Curracurrang and Nullarbor clusters. Panels **E, F**, and **G** show the distribution of the West, North, and East ancestral components, respectively. Colour transparency reflects the magnitude of the ancestry component geographically. Interpolation in **A, E, F**, and **G** are performed following **SI Section 4**.

To further resolve contemporary dingo population structure while minimising the confounding effects of European dog admixture, we analysed European dog ancestry-masked dingo genomes using principal component analysis (PCA) and ADMIXTURE clustering (see **SI Section 3.5, Fig. 2B, 2C, 2D**). Dimensionality reduction of the top 10 principal components using UMAP (McInnes et al. 2018) partitioned the individuals into eight genetically distinct clusters that correspond closely with their geographic provenance (**Fig. 2B, 2C**). Consistently, ADMIXTURE identified seven ancestral components as the best-supported model, which, when stratified by UMAP-defined groups, revealed distinct genetic signatures among populations (**Fig. 2D**).

Five of the eight inferred populations broadly correspond to clusters previously identified (i.e., West, East, Alpine, Mallee, and Captive) (Stephens et al. 2022; Cairns et al. 2023; Scarsbrook et al. 2025; Weeks et al. 2025). In support of our *f*_*4*_ analyses, individuals that show greater affinity to the Nullarbor lineage fall within the previously identified West group, whereas the East cluster encompasses the ancient Curracurrang dingoes and related modern individuals— though our results imply that this cluster is more extensive, spanning the majority of the Great Dividing Range (**Fig. 2C, 2E, 2G, S16**), with affinity to the Curracurrang lineage decreasing at higher latitudes. This latitudinal gradient coincides with a reduction in the eastern ancestral component inferred by ADMIXTURE (**Fig. 2E, S17**).

K’gari dingoes also form a distinct genetic cluster, consistent with small population size and long-term isolation from the mainland (Miller et al. 2024; Souilmi et al. 2024). We further identify two previously undescribed clusters, North and Central groups. The North cluster is concentrated in northern and central mainland Australia (**Fig. 2F**), and shows reduced affinity to the Nullarbor lineage compared to the West cluster (**Fig. 2A, S16**). The Central cluster comprises individuals from central Australia, and showing mixed ancestry from the North, East, and West clusters, consistent with their geographic position at the intersection of these groups (**Fig. 2C, S16**). Additionally, the population structure we describe can be broadly characterised by differential affinity to these ancient lineages—western populations predominantly share affinity with the Nullarbor lineage, eastern populations with the Curracurrang lineage, and central populations show no strong affinity to either (**Fig. 2A, 2C, S16**). Instead, central Australia appears to represent a zone of mixed ancestral dingo ancestry, and this pattern is further supported by elevated genetic diversity observed in these populations (**Fig. 3D**).

**Figure 3.**
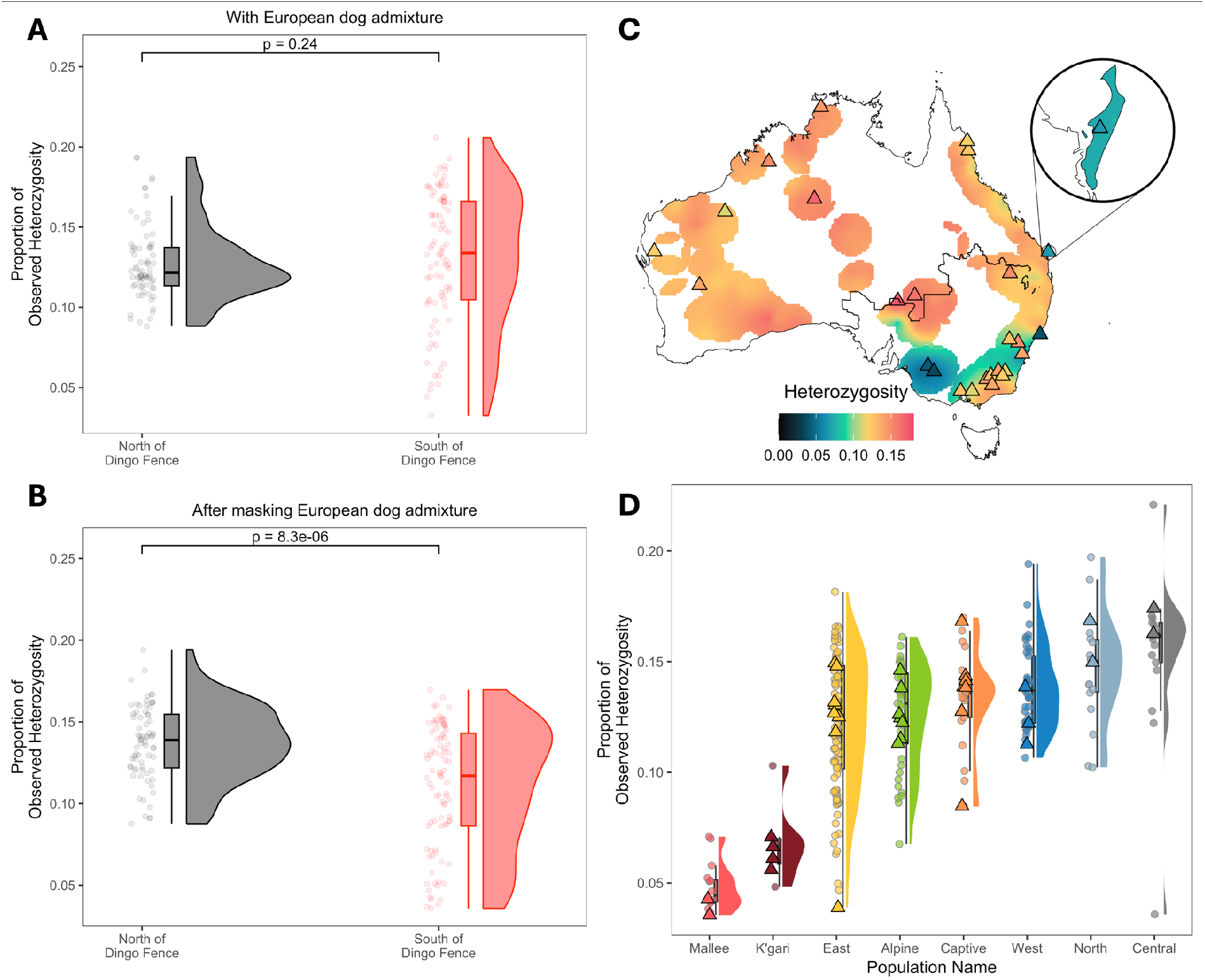
Ancestral Dingo Genetic Diversity. **A)** Distribution of dingo genetic diversity (calculated as a proportion of heterozygous alleles across the SNP-Array per individual) across the whole-genome north and south of the dingo fence. Plot is annotated with the p-value. **B)** As in panel **A**, but showing the distribution of dingo genetic diversity after masking European dog ancestry. **C)** Geographic distribution of ancestral dingo genetic diversity. Triangles represent whole-genome samples. **D)** Distribution of ancestral dingo genetic diversity stratified by populations defined in **Figure 2**.

### Ancestral dingo genetic diversity

A recent study suggested that admixture from European dogs may have partially alleviated inbreeding depression in southeastern populations subjected to intensive lethal control (Scarsbrook et al. 2025). To test this hypothesis, we compared dingo genetic diversity estimates with and without masking European dog ancestry, contrasting groups on either side of the dingo-proof fence (**Fig. 3A, 3B**). Here, we use the dingo fence as a proxy for contrasting land-use regimes in southeastern Australia. When European dog ancestry was included, no significant differences in genetic diversity were observed between populations north and south of the dingo fence (**Fig. 3A**). In contrast, masking European dog ancestry revealed significantly lower ancestral dingo genetic diversity in populations south of the fence compared to those north of the fence (**Fig. 3B**).

We next assessed dingo-specific genetic diversity across the eight genetic sub-groups identified in our previous analyses. Dingo-specific genetic diversity is highest in the Central Australian group, consistent with our ADMIXTURE results and the absence of strong affinity to either the Curracurrang or Nullarbor ancient lineages, suggesting that this region represents a zone of high gene flow relative to peripheral regions (**Fig. 3D, S16**). K’gari dingoes show low levels of genetic diversity, consistent with previous reports of a small population size (Miller et al. 2024). Notably, groups concentrated within southeastern Australia—i.e. Mallee, East, and Alpine dingoes—also exhibit markedly reduced ancestral diversity (**Fig. 3C, 3D**), with the Mallee group having even lower diversity levels than that observed on K’gari.

Spatial interpolation of ancestral dingo diversity further identified localised regions of reduced diversity within the East cluster (**Fig. 3C**). These putative genetic isolates are supported by spatio-genetic modelling using FEEMs (Marcus et al. 2021), which reveals significantly reduced effective migration rates in these regions (**Fig. S12**). Together, these patterns suggest that historical and recent anthropogenic landscape modifications may have reinforced isolation and reduced genetic diversity in southeastern dingo populations.

## Discussion

In this study, we demonstrate that estimates of European dog admixture in dingoes are highly sensitive both to the type of genetic assay used and the statistical inference method employed. Compared to qpAdm based estimates, STR data consistently overestimate European dog admixture across the species’ range (**Fig. 1A**), whereas clustering-based estimates from SNP-array and WGS data significantly underestimate European dog ancestry (**Fig. 1A**). Accurate and robust estimates of both historic and recent European dog ancestry in dingo populations is critical for dingo conservation programs. Imprecise ancestry estimates may lead to the inadvertent loss of ancestral dingo diversity through culling of individuals carrying high levels of dingo ancestry or the translocation of admixed individuals.

Our study demonstrates the formative role of natural biogeographic barriers in shaping contemporary dingo population structure. While these effects have been suggested in earlier studies (Cairns et al. 2023; Stephens et al. 2023; Weeks et al. 2025), they are more clearly resolved here, based on inferences applied to nuclear data that explicitly account for European dog ancestry. Our results confirm that modern dingoes retain deep affinities with ancient lineages represented by the Nullarbor and Curracurrang palaeogenomes, revealing population divisions that extend back thousands of years that were shaped by major biogeographic barriers. This is consistent with the relatively simple phylogeographic patterns inferred from mitochondrial DNA, where two divergent maternal lineages imply a longstanding east-west division either created by two distinct arrival events or an early post-arrival divergence (Cairns & Wilton 2016; Souilmi et al. 2024; Scarsbrook et al. 2025). Additionally, our nuclear genomic analyses reveal population subdivisions that likely reflect subsequent mixing and regional differentiation following the isolation of these lineages.

We demonstrate significantly lower ancestral genetic diversity in southeastern populations after masking European dog admixture (**Fig. 3A, 3B**). This pattern is consistent with recent findings that European dog admixture may have partially alleviated inbreeding depression in these populations (Scarsbrook et al. 2025). However, this apparent genetic rescue reflects admixture-driven increases in heterozygosity rather than the preservation of ancestral dingo diversity, and continued lethal control may further erode the ancestral genetic diversity. The Mallee population shows low genetic diversity with and without European dog alleles, likely a combined effect of historical and recent reductions in population sizes (Cairns et al. 2023; Scarsbrook et al. 2025). Interestingly, the Mallee and Central populations show comparable ancestral affinities and geographic proximity, with the Central population exhibiting the lowest genetic differentiation (F_ST_) to Mallee compared to all other populations (**Fig. S16, S18**). Together, this suggests historical connectivity between these regions and indicates that the Central population may represent a genetically compatible source that could help increase dingo specific genetic diversity in Mallee.

Levels of European dog admixture appear to be correlated with human population density (**Fig. 1B, 1C**). Despite our results suggesting ubiquitous admixture in southeastern Australia, previous studies have failed to find European dog mitochondrial lineages in dingo populations (Savolainen et al. 2004; Cairns & Wilton 2016; Scarsbrook et al. 2025). Indeed, discordance between nuclear and mitochondrial lineages are common in mammals, and could indicate sex-biased drivers of gene flow (Toews & Brelsford 2012). Furthermore, we confirm that the timing of admixture with European dogs is primarily historical (Scarsbrook et al. 2025), dating to around the 1950s, interestingly coinciding with the onset of widespread lethal control measures (van Eeden et al. 2019), and following a rapid post-World War II agricultural expansion of people—particularly in the southeastern regions (Godden 1999; Australian Bureau of Statistics 2024). Together, these interlinked factors may have resulted in reduced dingo population sizes, increased abundances of European dogs, and introduced constraints to dingo movement, limiting access to suitable conspecific mates. This combination provides a striking example of how anthropogenic pressures can rapidly and persistently reshape the genomic landscape of wild carnivores.

### Implications for future Dingo management and conservation

Given the potential for misclassification and mismanagement under the current genetic assays and associated methodologies, we recommend that conservation agencies adopt the gold standard testing methods, and continue updating guidelines as methods evolve and new data reference datasets become available.

While WGS remains the gold standard for population genetic inference, it has not been implemented in routine ancestry testing in conservation and management contexts due to financial and logistic barriers (Bertola et al. 2024). Our study demonstrates that qpAdm-based modelling, using ancient dingo genomic data as unadmixed sources, offers a more robust, scalable, and accurate method for ancestry estimation that remains reliable even with moderate amounts of pre-ascertained SNP markers, and as few as 10 thousand pseudohaploid transversions that are heterozygous in an outgroup (e.g., Coyote). The latter highlights the potential of low-pass shotgun sequencing as a cost-effective and practical approach for large-scale ancestry screening, reducing the need for high-coverage whole-genome and avoiding potential biases from pre-ascertained marker panels. Additionally, local ancestry inference approaches (e.g., MOSAIC (Salter-Townshend & Myers 2019)) provide another alternative to accurately account for European dog admixture when assessing ancestral dingo diversity and identifying critical conservation units and their historical relationships.

Our findings imply that the vast majority of modern dingoes only carry a small fraction of European dog ancestry, suggesting the usage of the term ‘wild dog’ is unlikely to be appropriate for most free-living canine populations in Australia. Moreover, recent evidence suggests that European dog admixture may improve adaptive potential and help mitigate effects of historical population decline (Scarsbrook et al. 2025). However, ongoing lethal control in southeastern Australia may continue to erode ancestral dingo diversity, likely eliminating important local adaptations that emerged over thousands of generations. These nuances highlight the need for research-informed management strategies that account for both the risks and potential ecological benefits of European dog admixture, alongside the long-term and recent genetic history of Australian dingoes.

## Conclusion

European dog admixture in free-roaming dingo populations is limited, largely historical, and associated with proximity to large population centres and past lethal control. Future management should focus on improving ancestral dingo genetic diversity in populations within the dingo fence and maintaining sufficiently large populations to prevent genomic erosion. We also advocate that future research, planning and conservation action should engage with local Indigenous Australian groups to produce strategies that are scientifically robust, ethically sound, and culturally appropriate. Finally, palaeogenomes provide a valuable baseline for assessing the impact of gene flow from recently introduced domesticates and guiding informed conservation action.

## Materials and Methods

### Genomic Data

We assembled a broad genomic dataset comprising previously published pre-colonical dingoes, present-day free-living and captive dingoes, and ancient and modern dogs from Eurasia, along with other extant canid species. Full details of data sources and processing steps are provided in **Supplementary Methods SI Section 1**.

### Proposed genomic ancestry testing

We use the qpAdm function (Harney et al. 2021) from the ADMIXTOOLS2 (2.0.0) (Maier et al. 2023) R package to estimate the genetic proportion of European dog and dingo ancestries, contributing to modern dingoes. The European dog ancestry was modelled using German Shepherd dogs (4 high-coverage samples) (Plassais et al. 2019). The dingo ancestry was modelled using the two ancient lineages, the Nullarbor cluster and Curracurrang cluster (Souilmi et al. 2024). Although, it should be noted that the individuals from the Curracurrang cluster were enriched using whole-genome enrichment before sequencing. As a result, we evaluate any bias that could arise from its use and find no bias when using both the ancient dingo lineages (see **SI Section 5.2**). A rotating qpAdm approach, as previously described (Bergström et al. 2020), was implemented where every combination of the three populations was used as sources to model a given dingo target sample. If any of the three populations were not used as the source for a given model, then it was used as a reference population. qpAdm was run with default parameters, except with ‘*allsnps*’ enabled as stated below.

Additionally, we used the “CoyoteCalifornia” (Coyote01), “Zhokhov9500BP” (CGG6), “Germany_7k” (Herxheim), “Ireland_Neolithic” (Newgrange), “Russia_Baikal_7k” (OL4223, C26 and C27) samples as constant reference populations, to represent global diversity outside of dingo and European dogs (see **Table S1**). We chose ancient dog samples that have previously been shown to capture diversity that is ancestral to most domestic dogs today, but which can not be confounded by recent admixture from modern European dogs (Bergström et al. 2020, 2022). Although free-living dogs from around the world are good candidates for the reference populations, many of them have been shown to be admixed with European breeds (Bergström et al. 2020), which can confound the models being tested here. The rotating qpAdm approach was run on dingo samples on a subset of 1.9 million biallelic transversions that are heterozygous in the outgroup “CoyoteCalifornia” from “Canids 19M Tvs” call-set and for dingo samples in call-set “Canids 46K Tvs”. qpAdm was run with “allsnps” set to true for call-set “Canids 46K Tvs” but not for call-set “Canids 19M Tvs”.

Estimated qpAdm models are considered viable if their *p*-values are ≥ 0.01 and if their admixture proportions are feasible; i.e. all admixture proportions are between 0 and 1. For samples where multiple models were viable, models with fewer source populations and low standard error for admixture proportions (Z ≥ 3) were preferred. For any other ties, models with the highest *p*-value were chosen.

## Acknowledgments

We acknowledge the traditional custodians of the lands from which specimens used in this study have been collected and express our deep respect and gratitude to their Elders and traditional owners. We extend our gratitude to all the colleagues that produced and made available the various datasets we used in this study. We are grateful to Dr. Peter J. S. Fleming (NSW Department of Primary Industries and Regional Development) and Greg Mifsud (Centre for Invasive Species Solutions) for their valuable feedback on the manuscript. Their insights helped refine the discussion and added important nuance and perspectives to a complex and rapidly developing topic. This work was supported by funding from the Environment Institute, Adelaide University, The Australian Research Council Centre of Excellence for Australian Biodiversity and Heritage (ARC CE170100015).

