## Supplementary Methods for "Palaeogenomics-informed inferences of European dog admixture enables scalable dingo conservation"

**Author Contributions:** B.L. and Y.S. conceived this study. S.R., N.C.N., L.T. and N.M.N. analysed the data. S.R., N.C.N., Y.S. and B.L. wrote the manuscript. All co-authors were involved in the interpretation of results and editing of the manuscript.

**Conflict of interest disclosure:** Y.S. receives funding from the Centre for Invasive Species Solutions, and the Department of Primary Industries and Regions, Government of South Australia.

**Keywords:** admixture, ancient DNA, conservation, dingo, genomics, heritage, palaeogenomics

### Supplementary Methods

#### SI Section 1: Genomic data

We acquired ancient whole-genomic data representing dingoes' ancestral heritage (n=14) (Souilmi et al. 2024; Scarsbrook et al. 2025). Particularly samples A19054 and A19058, (Souilmi et al. 2024) and 9 ancient dingoes (Scarsbrook et al. 2025) from Nullarbor, representing the Western dingo lineage, and samples D10, D13 and D16, from Curracurrang representing the Eastern dingo lineage. We also included 41 complete dingo genomes from around mainland Australia (Zhang et al. 2020; Scarsbrook et al. 2025), and supplemented this dataset with three K'gari samples (Souilmi et al. 2024) and Illumina shotgun-sequencing of the alpine (Ballard et al. 2023) and desert (Field et al. 2022) dingo reference genomes (**Fig. S1**), totalling 46 whole-genome dingoes. Additionally, we also included dingoes (n=391) and dogs (n=152) genotyped using the Axiom Canine HD Genotyping array (Thermo Fisher Scientific Inc.) (Cairns et al. 2023) (**Fig. S1**). Finally, we added 16 genomes from ancient and modern canids (Frantz et al. 2016; Botigué et al. 2017; Plassais et al. 2019; Bergström et al. 2020; Sinding et al. 2020). Samples used and metadata are summarised in **Table S1**.

##### SI 1.1: Data processing

We re-processed the 48 modern dingo genomes using the nf-core/eager (v2.4.5) pipeline (Yates et al. 2021, 2022) with parameters tuned for modern DNA. First, we trimmed the poly-G tails at the read termini using fastp (Chen et al. 2018) (0.20.1) and removed Illumina adapters with AdapterRemoval (Schubert et al. 2016) (2.3.2). We mapped the reads to the CanFam3.1 reference genome using bwa mem (Li & Durbin 2009) (0.7.17), eliminating alignments with a mapping quality of less than 20. Then, we used Picard Tools' (https://broadinstitute.github.io/picard/) MarkDuplicates (2.26.0) to remove duplicate reads. We re-processed all ancient dingo genomes according to the methods described previously (Souilmi et al. 2024).

##### SI 1.2: Genotyping

###### SI 1.2.1: For genome wide ancestry estimation

We employed GATK HaplotypeCaller (McKenna et al. 2010) (v4.6.1.0) with the "--alleles" and "EMIT_VARIANTS_ONLY" options, focusing on 19 million biallelic transversions from (Plassais et al. 2019). This ensured consistency with the ancient genomes from (Bergström et al. 2020). Before merging into the final call set, we set genotypes to be missing if they had fewer than half or more than three times the genome-wide coverage for each sample. For the ancient dingo genomes from (Souilmi et al. 2024), we produced two pseudo-haploid call sets using Pileupcaller (sequenceTools 1.5.2), calling random haploid genotypes on the 19 million biallelic transversions and 190 thousand SNPs from the Axiom Canine HD Genotyping array (Thermo Fisher Scientific Inc., Waltham, USA).

In total, we produced three call sets: (1) ancient and modern whole-genome canids from (Frantz et al. 2016); (Botigué et al. 2017); (Plassais et al. 2019); (Sinding et al. 2020); (Bergström et al. 2020); (Zhang et al. 2020); (Bergström et al. 2022); (Field et al. 2022); (Ballard et al. 2023), (Souilmi et al. 2024) and (Scarsbrook et al. 2025), called on 19 million biallelic transversions (Canids 19M Tvs); (2) ancient and modern dingoes and modern dogs from (Zhang et al. 2020); (Field et al. 2022); (Ballard et al. 2023); (Cairns et al. 2023), (Souilmi et al. 2024) and (Scarsbrook et al. 2025), called on 190 thousand biallelic SNPs (Canis 190K); and (3) a merged call set producing 46 thousand biallelic transversions (intersection) from the above two call-sets (Canids 46K Tvs).

###### SI 1.2.2: For local ancestry inference

Ancient dingoes genomes above 0.8x coverage and modern whole-genome dingoes were imputed and phased using GLIMPSE (Rubinacci et al. 2021) v1.1.1 and a reference panel containing more than 1,701 samples as previously described (Bougiouri et al. 2025; Scarsbrook et al. 2025). We retained only imputed variants with a minor allele frequency >0.01 in the reference panel and INFO score >0.8 in the imputed samples. Finally, we thinned the imputed VCF using vcftools --thin 1000, removing SNPs within 1kb of each other, resulting in >1.5 million autosomal variants.

Similarly, the 391 dingoes genotyped using the Axiom Canine HD Genotyping array, were imputed and phased with BEAGLE (Browning et al. 2018, 2021) v5.5 using the same reference panel as used to impute and phase the whole-genome samples. We retained imputed variants with a minor allele frequency >0.01 in the reference panel as before and with derived $r^2$ >0.8. Finally, variants were thinned as above, resulting in >1.1 million autosomal variants.

#### SI Section 2: Dingo Ancestry Analysis

##### SI 2.1: Cluster-based admixture estimates

We also performed clustering-based admixture analyses to estimate the proportions of European dog and ancient dingo ancestries in modern dingoes. We ran the following for the call-set “Canis 190K”, with one individual from a pair of individuals that were related by second degree or higher removed (see **SI Section 3.1**). Samples used for cluster-based admixture estimates are highlighted in **Table S1**.

We ran ADMIXTURE (Alexander et al. 2009) (1.3.0) in unsupervised mode, modeling from K = 2 to 14, with ten replicates for each K value and the seed set to the current time. We also used ADMIXTURE’s cross-validation (CV) procedure to observe cross-validation errors in each K value.

##### *SI 2.2: Local Ancestry Inference*

Genome-wide local and global ancestry estimates were inferred for more than four hundred dingoes using MOSAIC (Salter-Townshend & Myers 2019) v1.5.0. As previously described in (Scarsbrook et al. 2025), we used a reference panel of European dogs (n=55; excluding Australian derived breeds) (Bougiouri et al. 2025) and pre-contact ancient dingoes (n=11) as reference groups (**Table S1**). MOSAIC was run with default parameters separately for the imputed whole-genome and SNP-array genotyped dingoes. Generation since admixture was inferred by calculating the correlation in copying at increasing genetic distance from the reference groups along the genome. Admixture timing was calculated assuming a generation time of 3 years.

#### *SI 2.3: Admixture $f_4$-statistics*

We estimated the presence of dog admixture in dingoes using D-statistics expressed as $f_4$*(CoyoteCalifornia, GermanShepherdDog; Y, X)* where *Y* is either the Nullarbor 1k or Curracurrang 2k ancient dingo lineages and *X* is test modern dingo individuals. A significantly ($Z \geq 3$) positive $f_4$-estimate indicates admixture between dog and the test sample. We calculated the $f_4$-statistics on the 'Canis 190K' callset. We compared these to the qpAdm estimates (**Fig. S13**).

#### SI 2.4: Spatial analysis of dingo ancestry

We used a spatial beta regression (Ferrari & Cribari-Neto 2004) generalised linear mixed modelling framework to examine whether dingo ancestry proportions are correlated to human population density and position relative to the dingo fence (north vs. south). We did this using the glmmTMB (Brooks et al. 2017) package in R. Mean human population density data from the Australian Bureau of Statistics (Australian Bureau of Statistics n.d.) was taken from circular spatial buffers of 400 $km^2$ around geolocations of individuals in the SNP-array dataset (Cairns et al. 2023). This buffer size was chosen to approximate an upper estimate of dingo home range sizes inferred from satellite tracking data (Newsome et al. 2015). To account for the highly skewed distribution of human population density, we applied a $log_{10}(x+1)$ transformation. We compared mean human population densities at sampled locations north vs. south of the fence to examine covariation using a *t*-test. To correct for spatial autocorrelation in ancestry between individuals, we included a Moran Eigenvector Map (MEM) variable in the model, which captures spatial structure in the data (Griffith et al. 2019). We also included a random intercept for position relative to the dingo fence to determine unmeasured spatial variability in dingo ancestry proportions between regions north and south of the fence.

#### SI 2.5: Subsampling WGS loci

For the 46 unrelated WGS dingo individuals, we generated ten independent pseudohaploid replicates per individual (i.e., for each genotype, one allele was randomly sampled and the genotype set to homozygous for that allele). Within each pseudohaploid replicate, we randomly subsampled *'n'* SNPs per individual where *n* = 1,000 (approx. number of transversions in the DArTSeq data), 5,000 (approx. number of loci covered by the DArTSeq dataset), 10,000, 50,000 (approx. number of transversions in the SNP-Array data) and 100,000 (approx. half the number of SNPs covered by the SNP-Array). In total, this resulted in 50 replicates per individual, differing in both pseudohaploid calls and retained loci, resulting in 2,300 replicates across all individuals. Ancestry estimation was then performed for each replicate using qpAdm, as described in the **Materials and Methods**.

### SI Section 3: Dingo Population Structure

#### SI 3.1: Kinship and duplicate detection

We used KING (Manichaikul et al. 2010) v2.2.7 to determine relatedness of all dingoes and dog samples in the Canis 190K callset. Using KING, we also confirmed individuals that were genotyped in (Cairns et al. 2023) and shotgun sequenced in (Scarsbrook et al. 2025) are indeed duplicates. For subsequent analysis, we removed one from each pair of related samples up to second degree related, preferring to keep the WGS in a related pair of WGS and SNP-Array individuals.

#### SI 3.2: Masking European Dog ancestry

MOSAIC outputs local ancestry as a grid of evenly distanced genomic segments. For SNP-Array and WGS dingoes, we used the *grid_to_pos* function in MOSAIC to translate the local ancestry proportions to the ~190 thousand SNPs targeted in the SNP-Array. For each sample, the dog allele likelihood (*DAL*) was calculated as the average likelihood of each allele originating from European dogs across all loci. Genotypes were masked if *DAL* > 0.2, using the *--zero-cluster* flag in PLINK to remove likely European dog alleles across all contemporary dingo populations.

Adaptive admixture from European dogs was not assessed in the SNP-array dingoes, as the combination of SNP-Array ascertained loci and imputation artificially increased the proportion of dog alleles (**Figure S8**).

#### SI 3.3: Outgroup $f_4$-statistics

We used D-statistics to calculate the affinity of modern dingoes to either the ancient Nullarbor 1k or Curracurrang 2k lineages. We calculate outgroup-$f_4$, expressed as $f_4(X, CoyoteCalifornia; Y, AndeanFox)$, where $X$ is test modern dingo individuals with European dog ancestry masked (**SI 3.2**) and $Y$ is either the Nullarbor 1k or Curracurrang 2k lineages. The outgroup-$f_4$, similar to the outgroup-$f_3$, calculates the shared drift between $X$ and $Y$ populations. To perform the calculations, we ran the qpdstat function from the ADMIXTOOLS2 (Maier et al. 2023) R package, with parameters “allsnp” and “f4mode” set to true. The outgroup-$f_4$ values were normalised to obtain a single ancestry affinity value, where negative or positive values show affinity towards the Nullarbor or Curracurrang lineages, respectively.

$$affinity = \frac{f_4(X, CoyoteCalifornia; Curracurrang, AndeanFox) - f_4(X, CoyoteCalifornia; Nullarbor, AndeanFox)}{f_4(X, CoyoteCalifornia; Curracurrang, AndeanFox) + f_4(X, CoyoteCalifornia; Nullarbor, AndeanFox)}$$

#### SI 3.4: Fast Estimation of Effective Migration Surfaces (FEEMS)

To infer gene flow and connectedness of modern dingo populations, we used FEEMS (Marcus et al. 2021) (1.0.0) to evaluate effective migration surfaces. We performed FEEMS on 219 contemporary dingoes from the SNP-array data and 78 thousand SNPs with no missingness, with latitude and longitude coordinates of samples, along with genotype data, as inputs. The spatial graph was fit on the data with a lambda factor of 2, with the covariance converging in 217 iterations.

##### SI 3.5: Principal Component Analysis (PCA) and ADMIXTURE

We performed PCA using EMU (Meisner et al. 2021) (v1.6.0), a software optimised for the presence of random and non-random missingness in genomic data, to account for the ancient samples and missingness introduced by masking European dog ancestry. EMU was run with default options. Only ancient and modern dingoes were included (**Table S1**), excluding individuals with less than 70% dingo ancestry, samples with fewer than expected alleles masked as European dog (**Fig. S14**), individuals with > 50% loci removed and one individual from each pair of related individuals. EMU generates output files containing the first 10 eigenvectors and eigenvalues, representing the variance in dingo population structure independent of European dog admixture. UMAP (McInnes et al. 2018) was then applied to these PCs to visualise distinct genetic clusters.

On the same set of individuals, we ran unsupervised ADMIXTURE following the same method detailed in **SI Section 2.1**.

#### SI Section 4: Spatial interpolation of dingo ancestry

Results of dingo ancestry estimates were obtained from qpAdm (see **Methods**), and affinity estimates (see **SI Section 3.3**) were mapped using Rnaturalearth *(South et al. 2025),* including kriging (gstat *(Gräler et al. 2016)*) analysis to interpolate to areas not sampled. For the SNP-Array and WGS data, one individual from a pair of related individuals was removed. The mean ancestry or affinity value was taken for locations with multiple samples. The grid to interpolate was created such that there were at least two samples within 200 km (an arbitrary choice for the sake of visualisation) of the interpolated point. A spherical model was fitted using the *fit.variogram* function. Samples included in the interpolation are highlighted in **Table S1**.

### Supplementary Results

#### SI Section 5: Evaluating different qpAdm models

To assess the robustness of the qpAdm models, we tested them across a range of variables to identify the most reliable applications and interpretations of the results.

##### SI Section 5.1: Accessing sources of European dog

We selected German Shepherd dogs as the source population to measure European dog gene flow, as most other European dog breeds cluster in a single clade with German Shepherds (Bergström et al. 2020). Nevertheless, we also evaluated the impact of source choice by using English Cocker Spaniels, Border Collies, Labrador Retrievers or Scottish Terriers instead (**Fig. S2**). There were no significant differences in ancestry estimates when using different European dog sources. We also observed that some individuals with low European dog ancestry (<10%) sometimes showed no detectable European dog contribution. The potential reasons for this are explored in **SI Section 5.3**.

##### SI Section 5.2: Accessing sources of ancient dingo

Two ancestral sources of dingo populations with genetic continuity to the present day were identified (Souilmi et al. 2024). However, the ancient individuals from the Curracurrang lineage were enriched using modern K'gari dingo genomic DNA during library preparation, potentially introducing bias. To evaluate how this might affect ancestry estimates, we ran models using only the Nullarbor lineage (shotgun sequenced without enrichment), only the Curracurrang lineage (shotgun sequenced with enrichment), or both lineages as sources (**Fig. S3**).

For WGS individuals, the mean difference in dingo ancestry estimates when using only Nullarbor or Curracurrang compared to both lineages was 0.0034 ± 0.023 and 0.0025 ± 0.023, respectively. For SNP-array individuals, however, using only Curracurrang as a source produced larger differences (0.037 ± 0.044) than using only Nullarbor (0.0089 ± 0.034). Because no such deviation was observed in WGS samples, this suggests the bias may be exacerbated by the combination of SNP-array ascertainment and enrichment of ancient Curracurrang samples. Nevertheless, including both lineages as sources appears to overcome this bias even in SNP-array samples, and we therefore include both lineages in the qpAdm models. Indeed, using only the Nullarbor lineage as a source would also be sufficient.

##### SI Section 5.3: Accessing the impact of reduced markers and haploid calls

To evaluate the effect of data missingness and pseudohaploid genotypes, we subsampled the WGS individuals as described in **SI Section 2.5**. We found that qpAdm provides robust ancestry estimates with as few as 10,000 haploid SNPs when using two-source models (**Fig. S4B**). However, the power to reject single-source models (i.e., models where individuals have only one source—ancient dingoes) decreases with fewer SNPs. In particular, individuals with lower dog ancestry require more SNPs to confidently reject single-source models (**Fig. S4A**).

##### SI Section 5.4: Assessing impact of SNP-Array Ascertainment

To assess the impact of SNP-Array ascertainment on qpAdm models, we downsampled whole-genome shotgun-sequenced dingoes to 1.2 million heterozygous transversions in the "CoyoteCalifornia" individual (Plassais et al. 2019) or to ~46 thousand transversions that overlap with the SNP-Array markers (Cairns et al. 2023) and compared the results (**Fig. S5B**). We found no systematic bias (mean difference between two sets of transversions = 0.004 ± 0.064), indicating that the SNP-Array loci are sufficiently informative to robustly differentiate between dingo and dog ancestries.

##### SI Section 5.5: Assessing impact of DArTSeq Ascertainment

We performed an analogous comparison for the five thousand DArTSeq loci (Weeks et al. 2025), this time downsampling the shotgun-sequenced dingoes to a set of approximately one thousand DArTSeq transversions. In contrast to the Axiome Canine HD Genotyping array, all analyses of the DArTSeq transversions returned qpAdm models with only an ancient dingo component, indicating that this assay has insufficient resolution to detect dog admixture when using such few markers (**Fig. S4**). This is consistent with our results from **SI Section 5.3** showing the need for at least 10,000 loci for accurate estimates.

Based on these results, we conclude that the qpAdm modelling framework proposed here is generally robust to different data types, sources of ancestry, and number and ploidy (i.e., diploid or haploid genotypes) of loci, in line with other benchmarking of qpAdm modelling (Harney et al. 2021). Additionally, the use of haploid SNPs, which is analogous to extreme homozygosity/inbreeding, indicates that this method is robust to low-population sizes. We advise caution when applying qpAdm to samples where both the source and target populations may have been ascertained using non-shotgun methods. Additionally, analyses with a reduced number of loci may have reduced power in differentiating between single- and multi-source models when the introgressed alleles are present in low quantities (i.e., lower European dog admixture in an individual).

#### SI Section 6: Comparing qpAdm to other ancestry estimation methods

##### SI Section 6.1: Comparing qpAdm to ADMIXTURE

To explore if the European dog admixture estimates are affected by choice of inference procedure, we compared our revised qpAdm-based estimates to those obtained by running ADMIXTURE (Alexander et al. 2009) on both WGS and SNP-Array datasets (see **Supplementary Methods SI Section 2.3**). ADMIXTURE is a widely used ancestry decomposition inference method that uses a clustering-based approach analogous to that adopted by FastStructure, the method that was employed to infer European dog admixture in the original dingo SNP-Array study (Cairns et al. 2023).

Consistent with previously reported FastStructure results, ADMIXTURE systematically overestimates dingo ancestry for the SNP-Array data, with similar overestimates also observed for WGS individuals (mean difference: 0.099 $\pm$ 0.11) (**Fig. S5C**). Notably, ADMIXTURE estimates of dog admixture in contemporary dingoes decreases as the assumed number of ancestral populations increases (**Fig. S10B**), especially in the “East” and “Alpine” subgroups. This suggests that clustering-based methods may be sensitive to post-admixture genetic drift, leading to inaccurate estimates of admixture proportions. Although, we observed no relationship between timing of admixture and discrepancy between qpAdm and ADMIXTURE estimates (**Fig. S9D**), indicating that differences between these methods are unlikely to be driven by temporal variation in admixture alone. These findings underscore the limitations of clustering-based methods when directly estimating admixture levels.

#### SI Section 6.2: Comparing qpAdm to MOSAIC

We next compared qpAdm ancestry estimates with those inferred by MOSAIC (Salter-Townshend & Myers 2019), a local-ancestry inference method recently applied to detect moderate European dog admixture in southeastern dingoes (Scarsbrook et al. 2025). For SNP-Array individuals, MOSAIC significantly overestimated European dog ancestry (p-value = 2.843e-09) relative to qpAdm by 0.046 ± 0.065 (**Fig. S8A**). The inflated European dog ancestry in the SNP-array data likely results from a combination of ascertainment bias, where the SNP-array targets variable loci in domestic dogs, and subsequent imputation using a reference panel predominantly representing domestic dog haplotypes. In contrast, WGS samples only show a slight overestimation of European dog ancestry by MOSAIC relative to qpAdm (0.012 ± 0.025; **Fig. S8B**), however this difference was not significant (p-value = 0.45) consistent with reduced ascertainment bias when imputing in shotgun-sequence data. Interestingly, nine individuals that were genotyped using both SNP-Array and whole-genome sequencing show broadly comparable estimates (**Fig. S7B**), indicating the combined ascertainment bias from SNP-Array and imputation do not affect all samples equally.

### Supplementary Figures

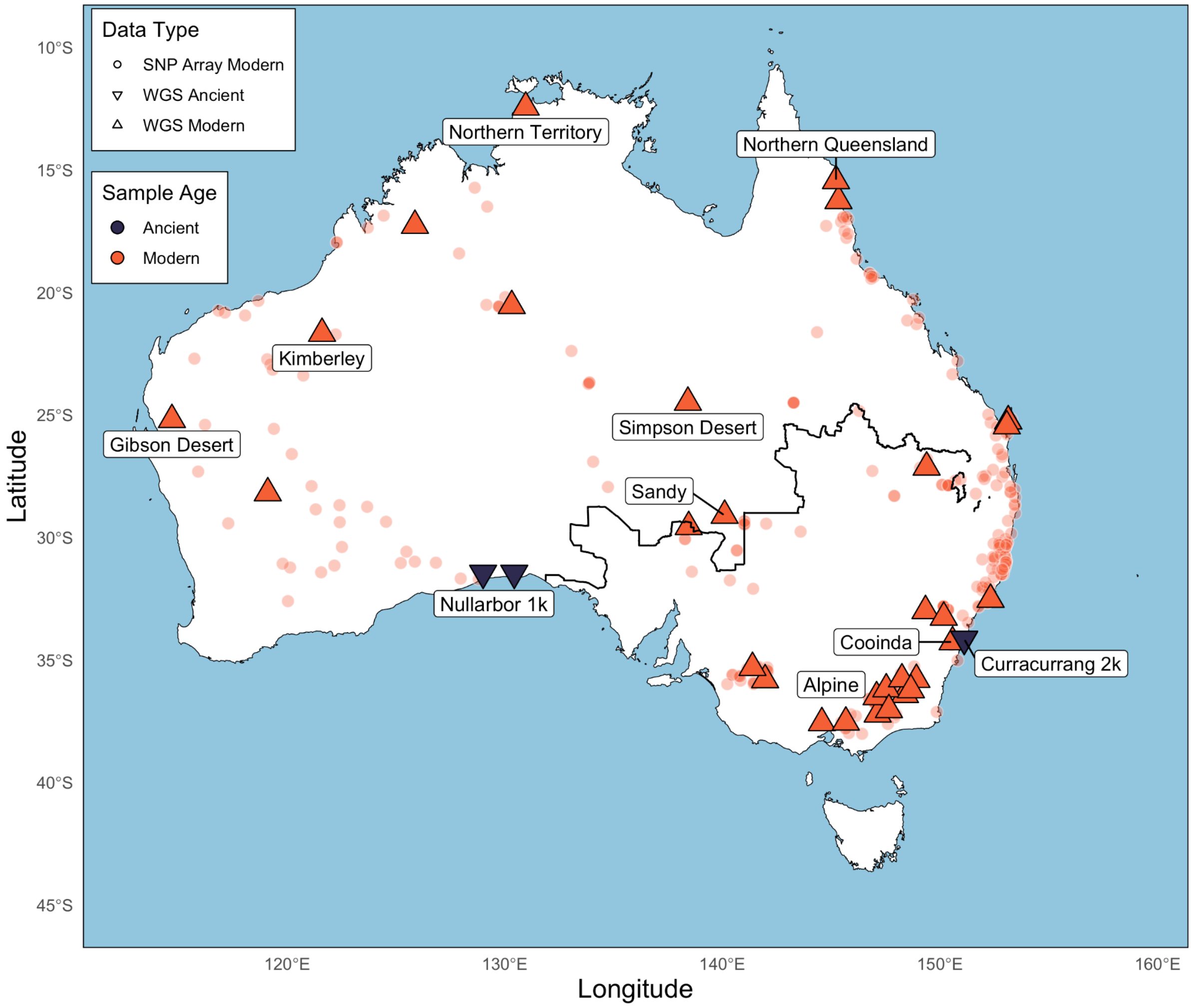


**Figure S1 | Sample Locations used in the study.** Sample locations of dingoes from (Ballard et al. 2023) (Cooinda reference genome), (Cairns et al. 2023), (Field et al. 2022) (Sandy reference genome), (Scarsbrook et al. 2025) (modern samples only), (Souilmi et al. 2024) and (Zhang et al. 2020). Translucent red circles indicate SNP-array data (Cairns et al. 2023). Opaque red triangles represent modern dingo whole-genome sequencing data (Field et al. 2022; Ballard et al. 2023; Souilmi et al. 2024), while opaque navy triangles denote ancient dingoes, including the Nullarbor 1k and Curracurrang 2k clusters (Souilmi et al. 2024). The map highlights the dingo fence in bold black line.

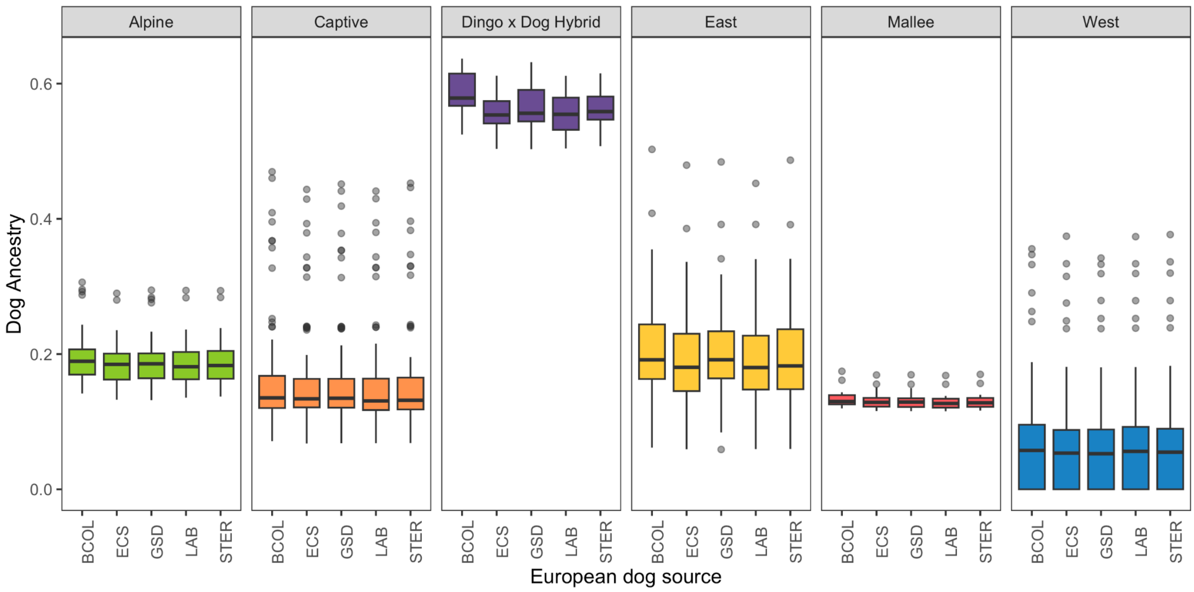


**Figure S2 | Comparing the effect of European dog source.** qpAdm estimates of European dog admixture levels in dingoes using different sources of European dog including Border Collie (BCOL), English Cocker Spaniel (ECS), German Shepherd Dog (GSD), Labrador Retriever (LAB) and Scottish Terrier (STER). The plot is faceted by populations. There were no significant differences.

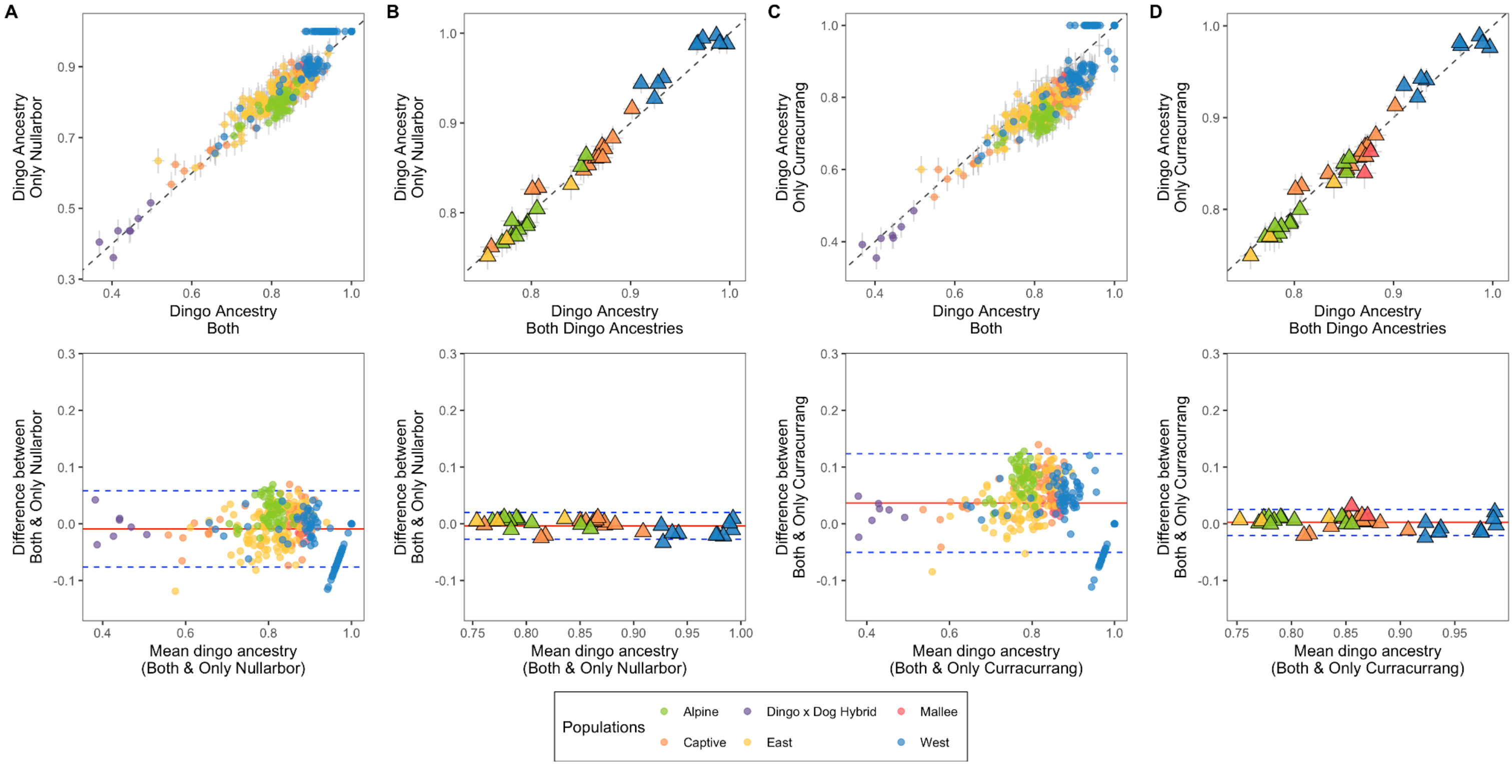


**Figure S3 | Comparing the effect of ancient dingo sources.** qpAdm estimates of dingo ancestry using both Nullarbor and Curracurrang lineages as sources, and using only either Nullarbor (panels **A** and **B**, comparing between SNP-Array and WGS individuals) or Curracurrang (panels **C** and **D**, comparing between SNP-Array and WGS individuals) as source. Each panel is divided into two sections: Top - Direct comparison between models using both ancient sources and those using a single source. Bottom - Bland-Altman plot, where the red line represents the mean difference in dingo ancestry estimates between models using both ancient lineages and those using a single lineage, and the dashed blue lines indicate ±1 standard deviation from the mean. Larger deviations from zero indicate greater differences between methods.

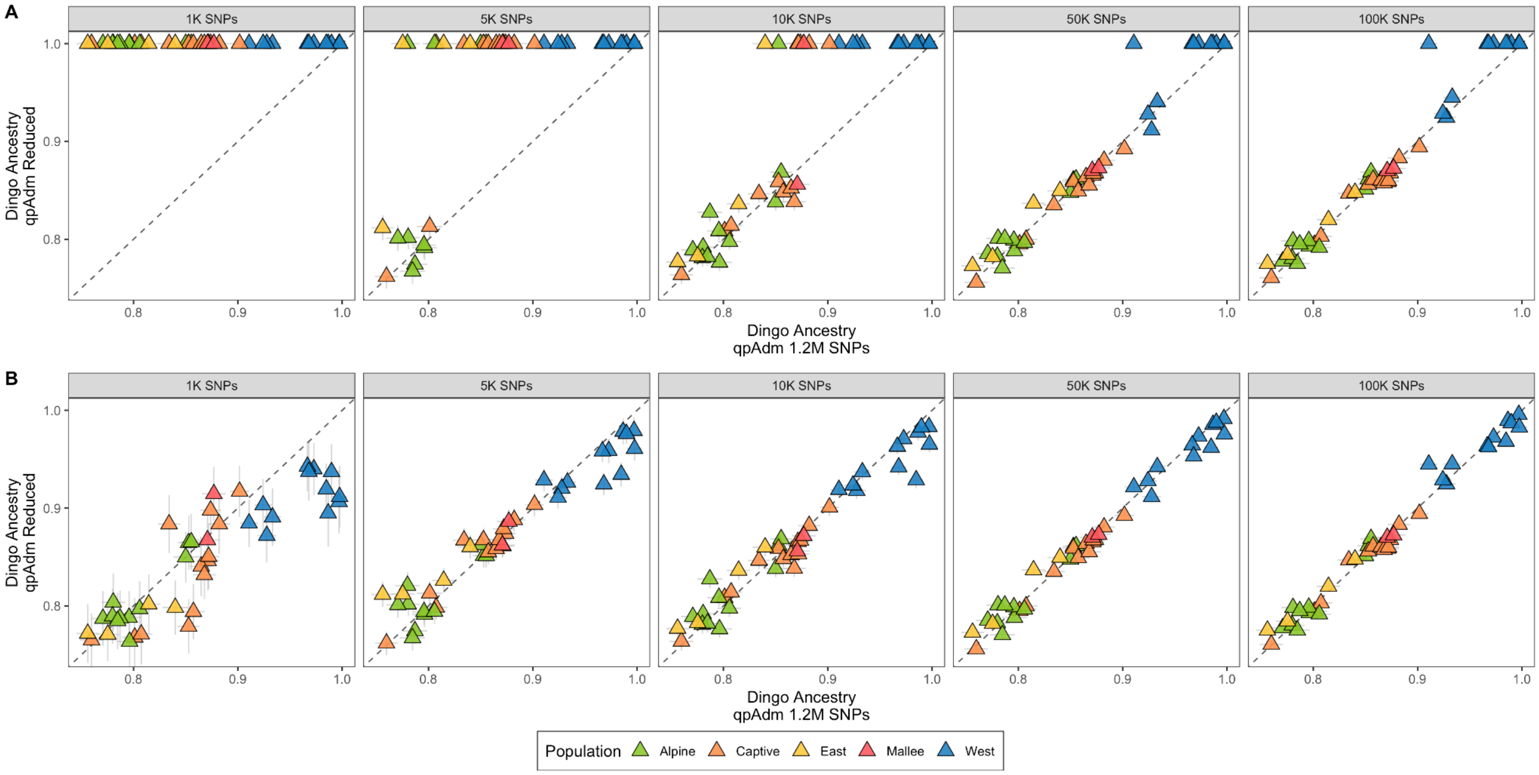


**Figure S4 | Comparing the effect of data quality.** qpAdm estimates comparing dingo ancestry estimates using 1.2 million transversions heterozygous in "CoyoteCalifornia" individual compared to reduced loci, as represented in the facets ranging from 1,000 to 100,000 pseudo-haplodised SNPs (see **SI Section 2.5**). Ten replicates were generated for each sample, varying both the pseudohaploid calls and the subset of SNPs used. Samples are coloured by population. **A)** Both single- and two-source models were considered, with preference given to models with fewer sources when multiple models were accepted. **B)** Only two-source models were considered.

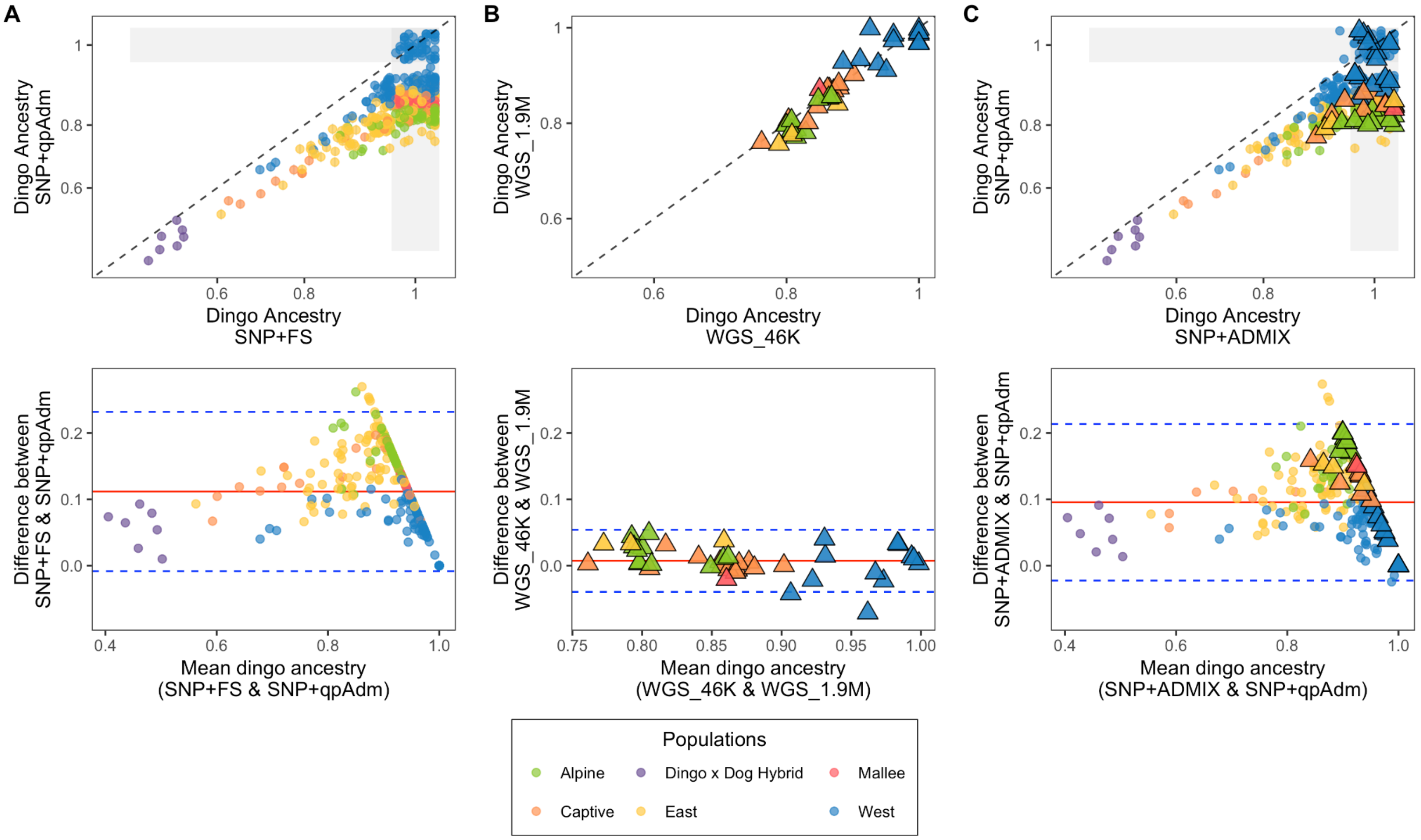


**Figure S5 | Comparing dingo ancestry estimates across different methods.**
**A)** Revised qpAdm estimates compared to previous FastStructure estimates. **B)** qpAdm estimates using two SNP sets: 1.9M transversions heterozygous in "CoyoteCalifornia" and 46K transversions from the SNP-array. **C)** Revised qpAdm estimates compared with the optimal ADMIXTURE k-value output. In all panels, samples are colored by population cluster. Circles represent SNP-array individuals, while triangles represent whole-genome sequenced individuals. Each panel is divided into two sections: Top - Direct comparison between the two methods. Bottom - Bland-Altman plot, where the red line represents the mean difference between methods, and the dashed blue lines indicate ±1 standard deviation from the mean. Larger deviations from zero indicate greater differences between methods. All points within the shaded area around 1 in plots **A** and **C** have an x and/or y value of 1.

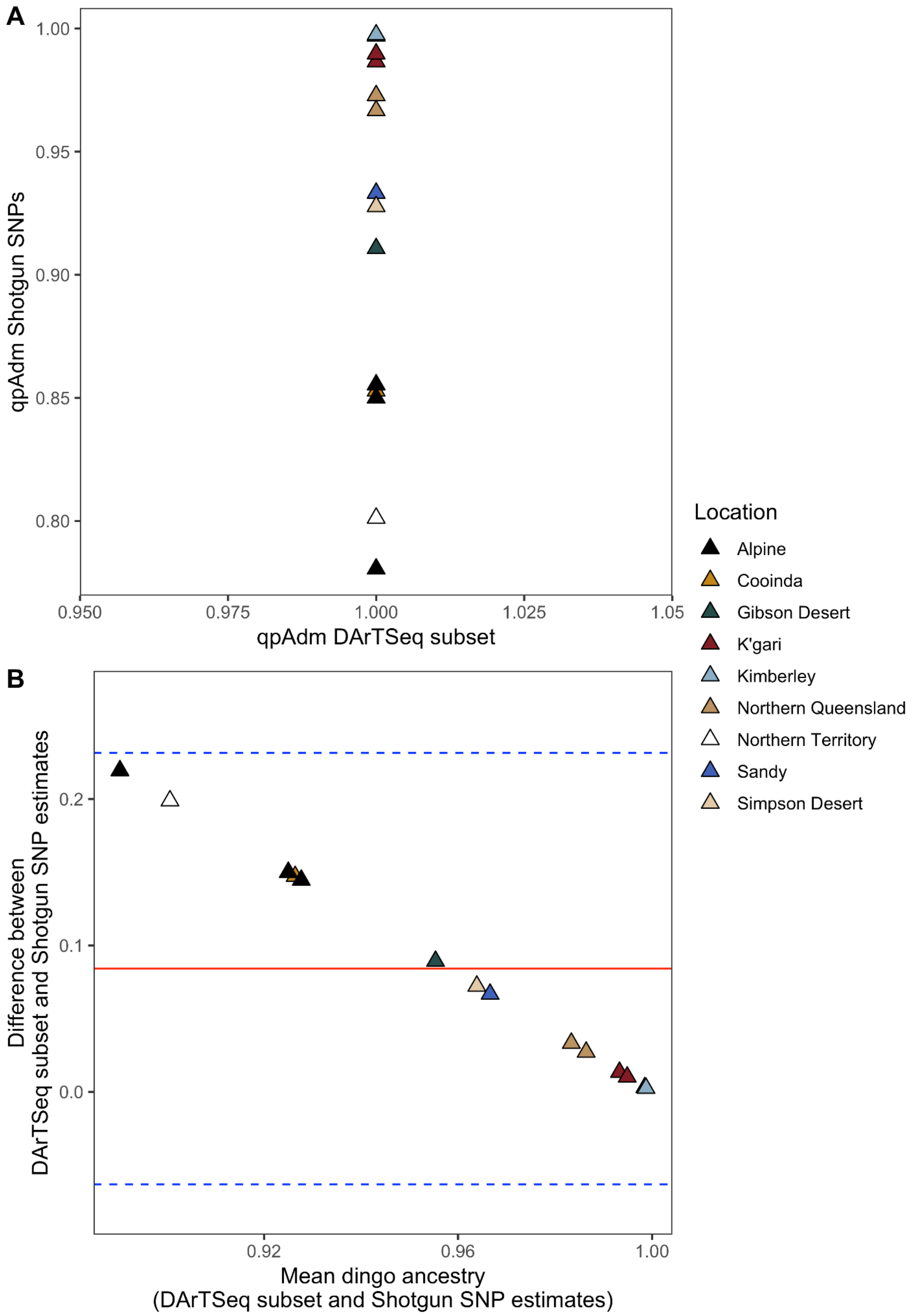


**Figure S6. A)** qpAdm models using 1k transversion from Weeks *et al.,* 2024 does not have enough resolution to differentiate dingo-dog admixture. **B)** Bland-Altman plot comparing difference and mean of qpAdm estimates using WGS and DArTSeq transversions.

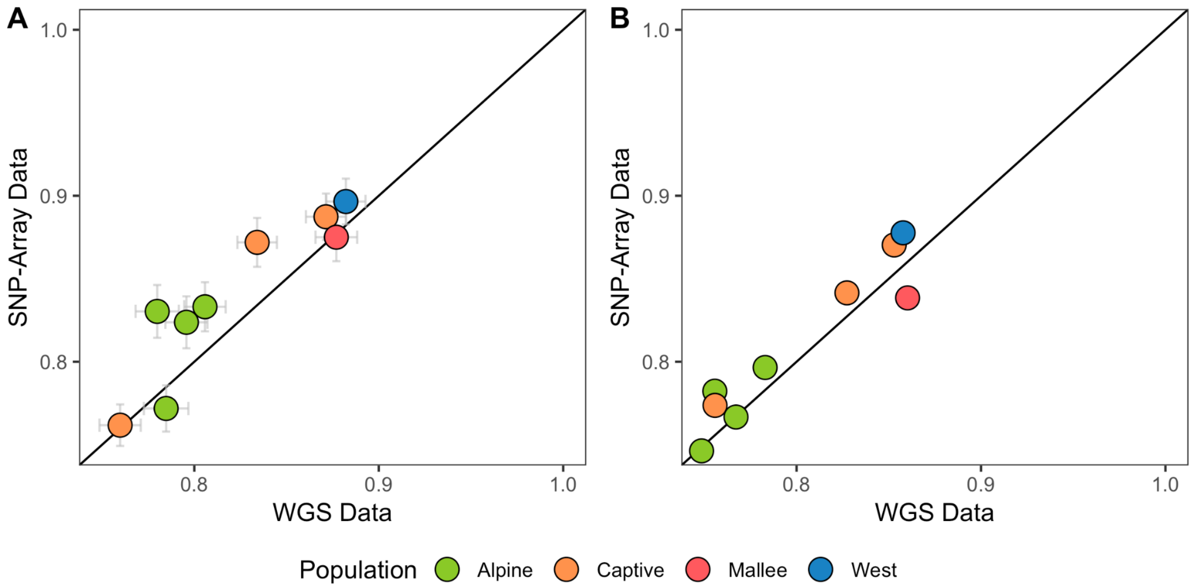


**Figure S7.** Dingo ancestry estimates of individuals that were genotyped using both SNP-Array and WGS using qpAdm (**A**) and MOSAIC (**B**).

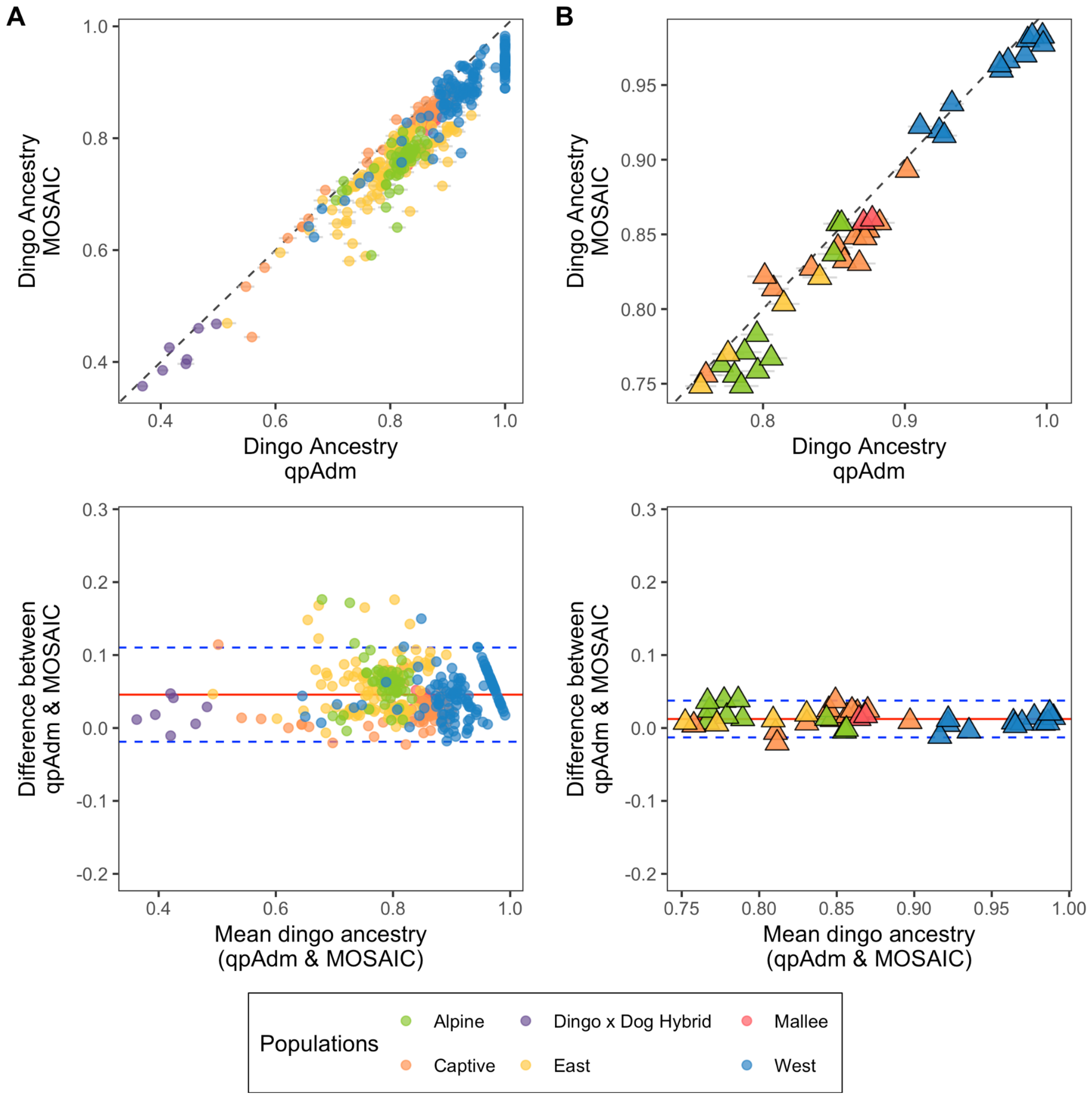


**Figure S8 | Local-Ancestry-Inferences (MOSAIC) vs. genome wide estimates (qpAdm).** Comparison of dingo ancestry estimates from MOSAIC and qpAdm for SNP-array and WGS samples (panels **A** and **B**, respectively). Each panel is divided into two sections: Top - Direct comparison between the two methods. Bottom - Bland-Altman plot, where the red line represents the mean difference between methods, and the dashed blue lines indicate ±1 standard deviation from the mean. Larger deviations from zero indicate greater differences between methods.

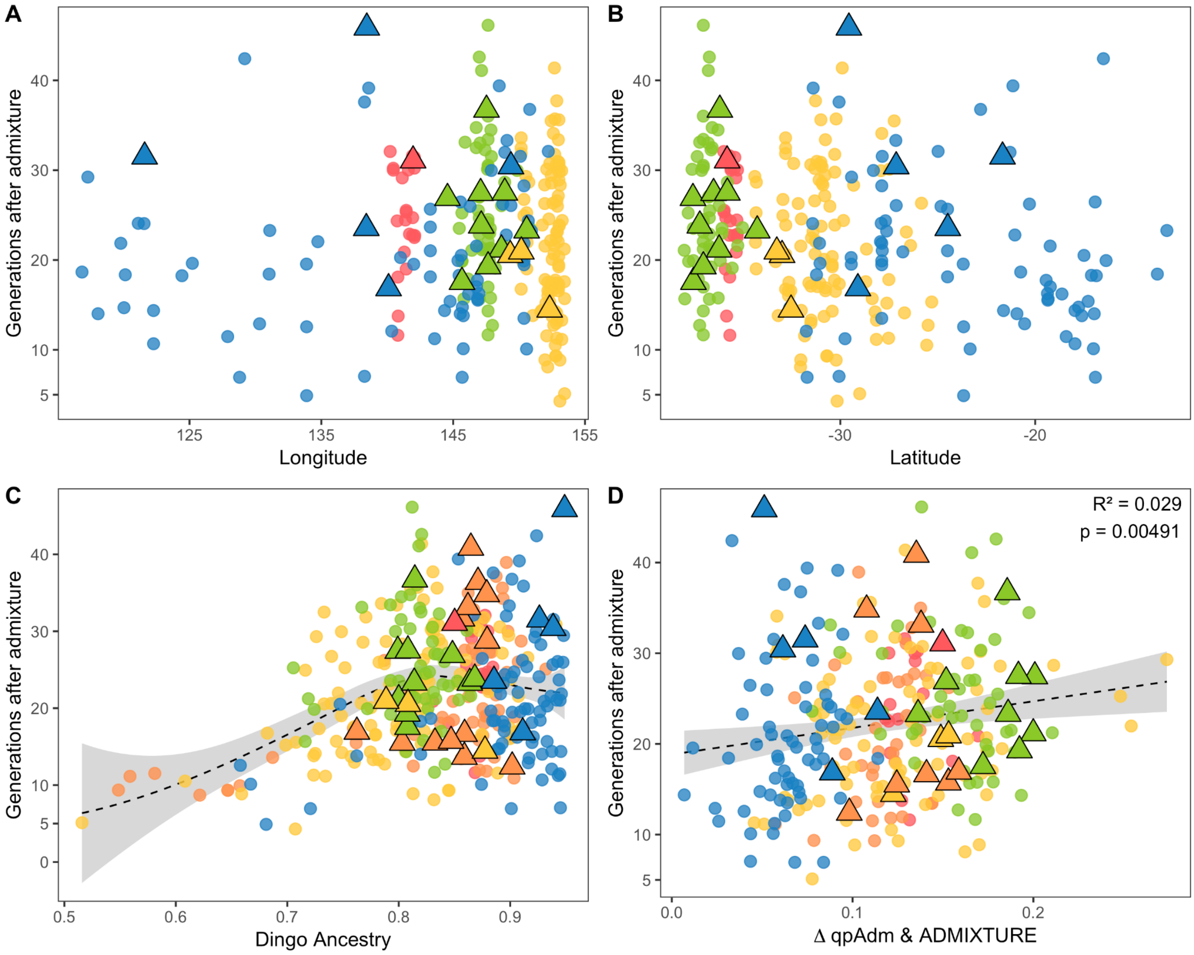


**Figure S9 | Timing of admixture with European dogs.** Admixture timing was estimated for samples with less than 95% dingo ancestry across longitude (**A**) and latitude (**B**). **C)** Relationship between dingo ancestry and admixture timing. **D)** Difference between qpAdm and ADMIXTURE estimates of dingo ancestry and admixture timing.

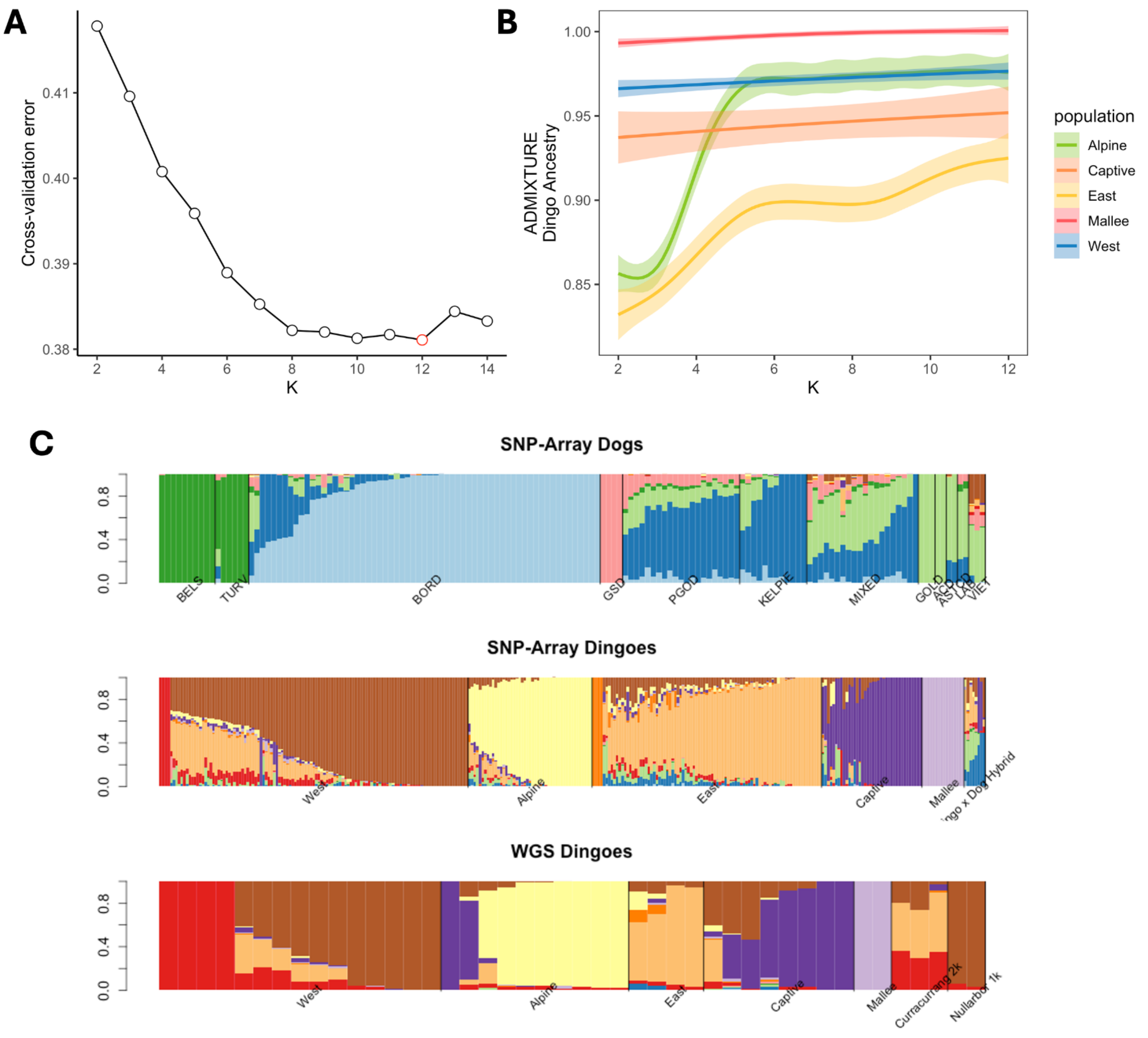


**Figure S10 | ADMIXTURE Summary. A)** Cross-validation (CV) error for ADMIXTURE run for K 2 to 14 on 505 ancient and modern dingoes and modern dogs. The K with the lowest CV error is highlighted in red. **B)** Dingo ancestry estimated using ADMIXTURE for captive and free-ranging dingoes at different K values (2 to 12). **C)** ADMIXTURE results for K=12. Populations include ancient dingoes from Souilmi *et al.,* 2024: Nullarbor 1k cluster and Curracurrang 2k cluster; WGS sequenced samples; samples from Cairns *et al.,* 2023: Mallee, Captive, East, Alpine, West, and Dingo x Dog Hybrid. Populations of dog used: Australian Cattle Dog (ACD), Australian Stumpy Tail Cattle Dog (ASTCD), Belgian Shepherd (BELS), Border Collie (BORD), Golden Retriever (GOLD), German Shepherd Dog (GSD), Kelpie (KELPIE), Labrador (LAB), Patagonian ovejero sheepdog (PGOD), Tervuren Shepherd (TURV), Vietnamese village dog (VIET), and Mixed breed (MIXED).

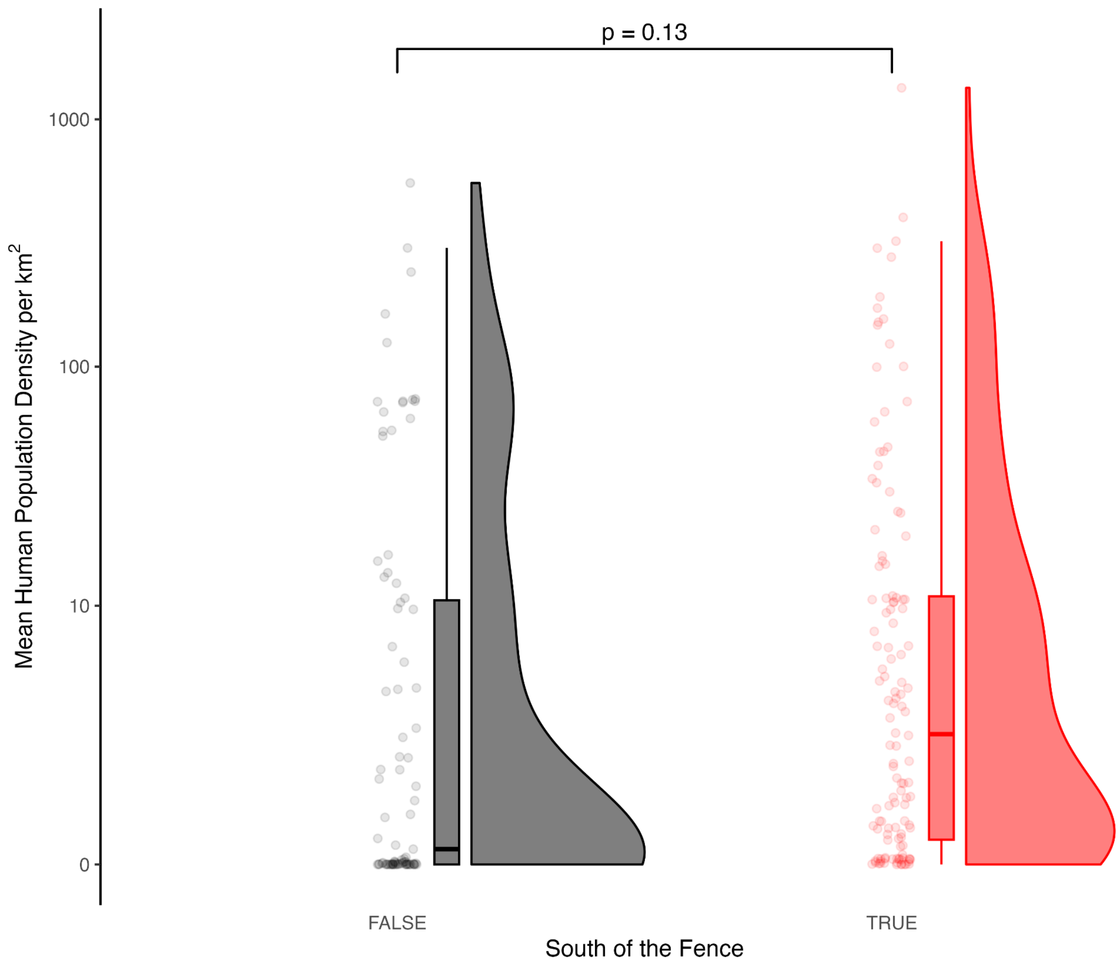


**Figure S11:** Mean human population density 400 km$^2$ around each dingo sample and position relative to the fence. There is no significant effect of relative position to the fence and mean human population density around dingoes. A standard t-test was performed here. The y-axis is on a log scale.

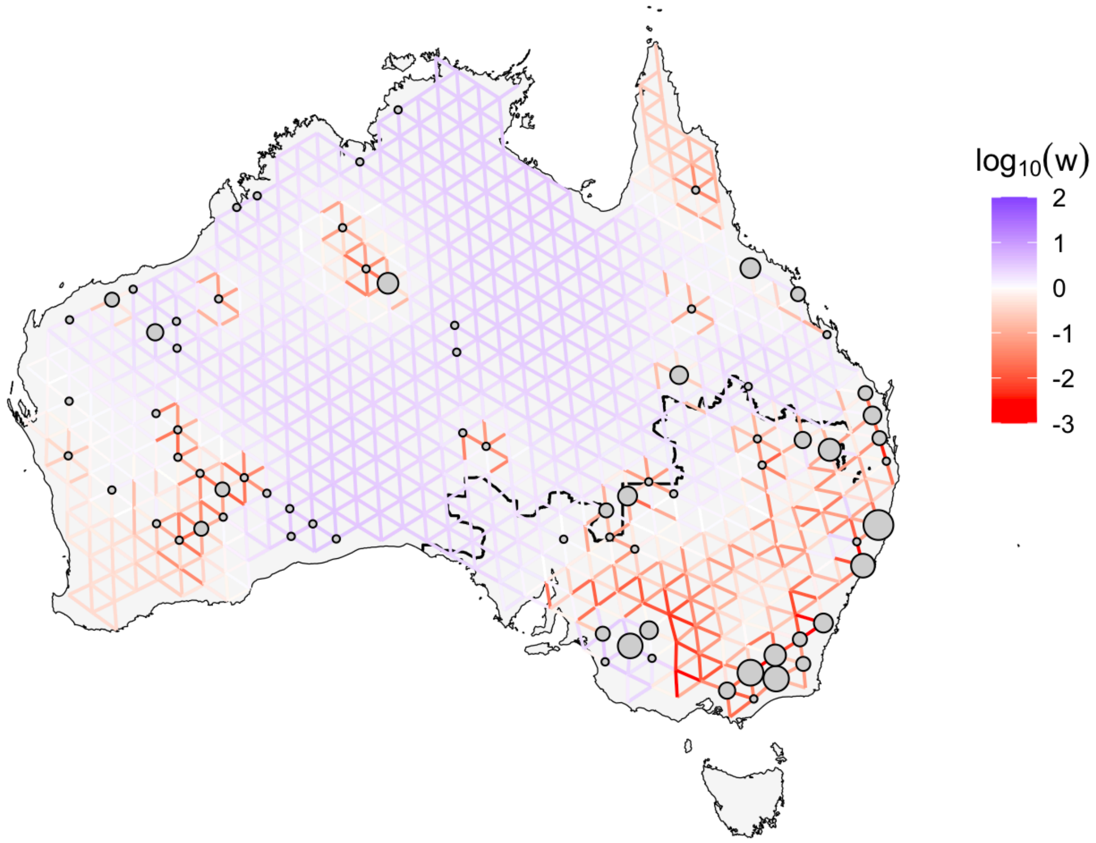


**Figure S12:** Effective Migration Surfaces of 219 contemporary dingoes from the SNP-array data and 78 thousand SNPs with no missingness. Positive values (blue) indicate higher than expected population connectivity and negative values (red) indicate lower than expected population connectivity.

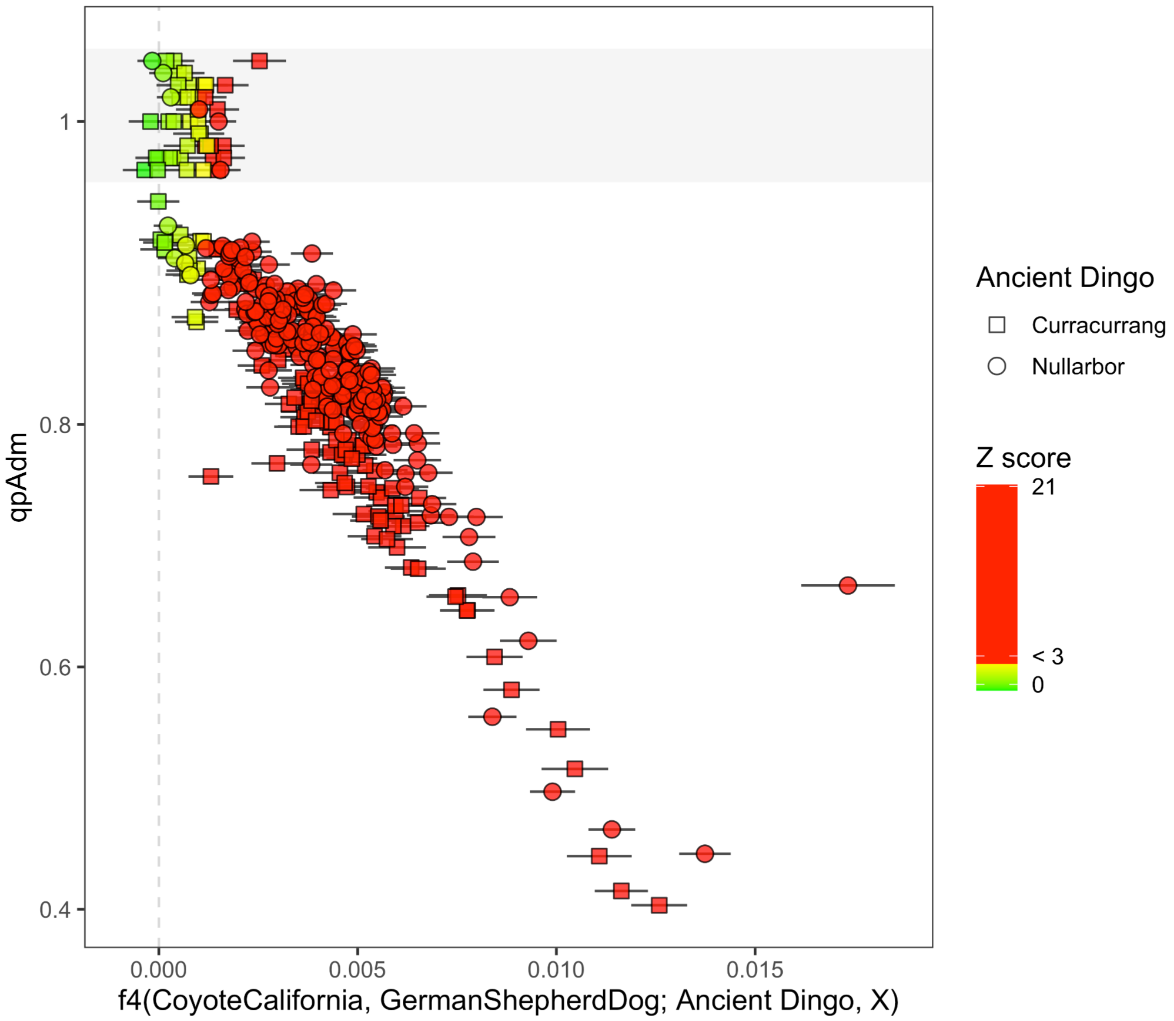


**Figure S13:** Comparing qpAdm estimates with $f_4$(*CoyoteCalifornia*, *GermanShepherdDog*; *Ancient Dingo*, *X*) where X is modern dingo samples. The points are coloured by Z-score and the shapes denote the ancient dingo source used to model the sample using qpAdm and calculate $f_4$ using all 190k SNPs. The shaded grey area at 1 all denote a qpAdm value of 1 (y-axis).

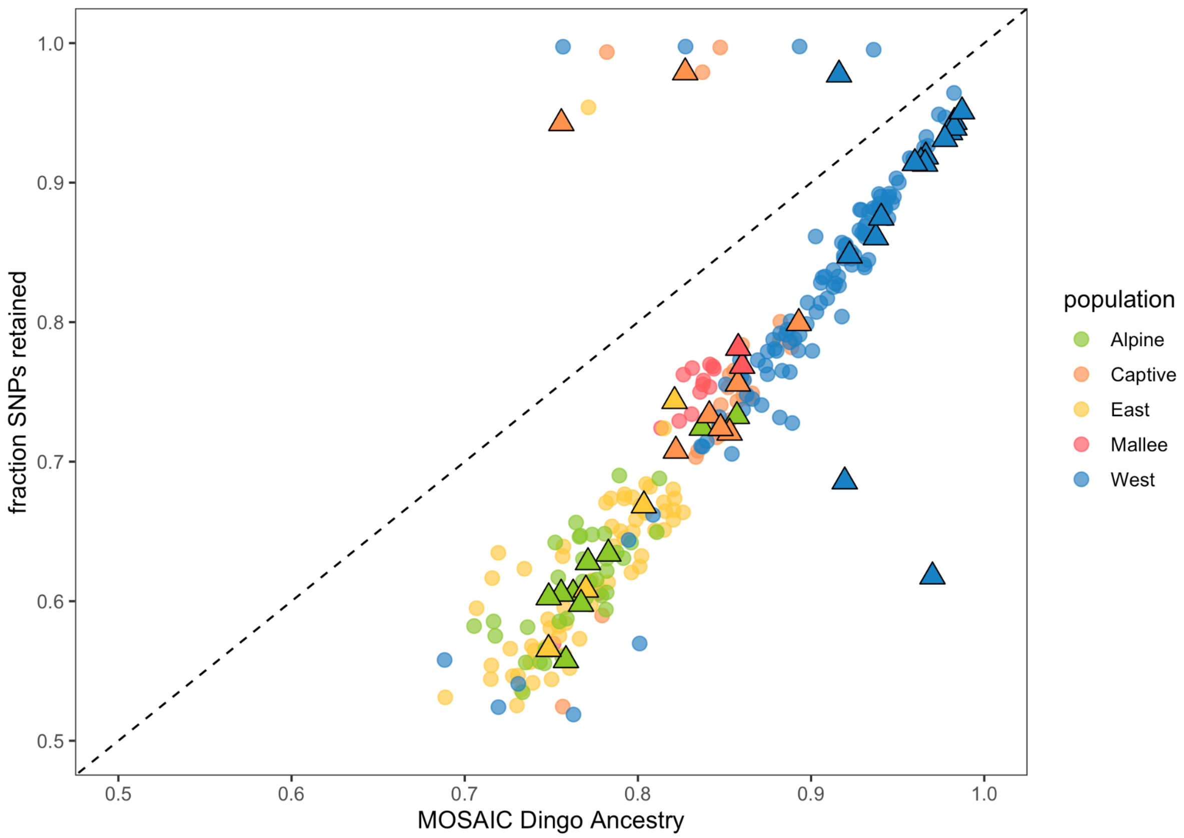


**Figure S14:** Relationship between MOSAIC Dingo ancestry estimates and fraction of SNPs retained after masking European dog ancestry. Samples above the $x = y$ line were removed.

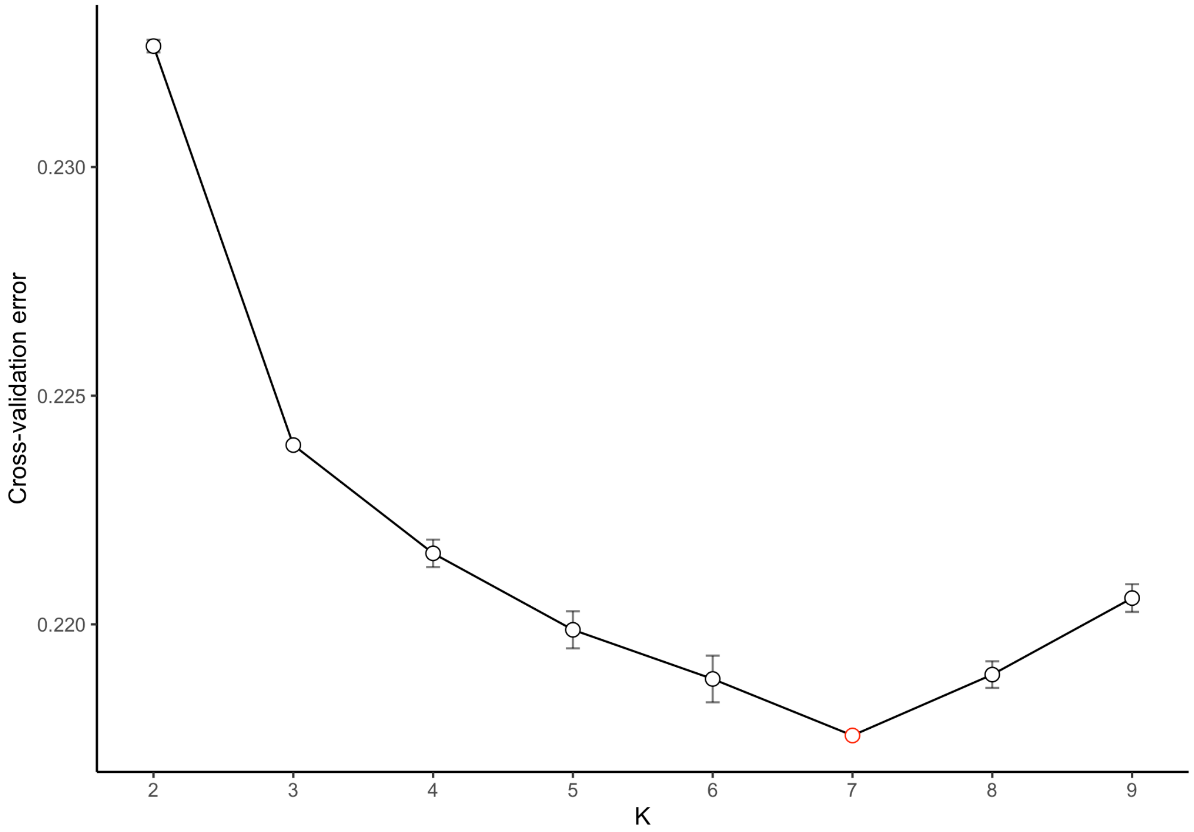

**Figure S15:** Cross-validation (CV) error for ADMIXTURE run for K 2 to 9 on 283 ancient and modern dingoes after masking European dog ancestry. The K with the lowest CV error is highlighted in red.

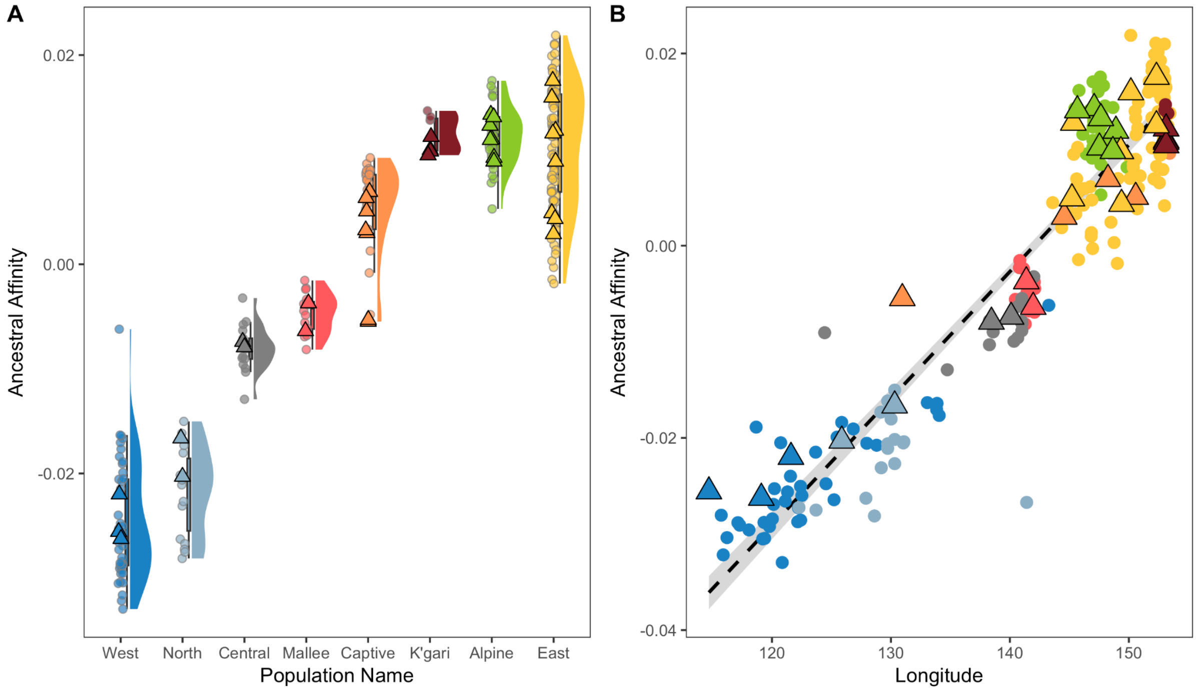


**Figure S16 | Ancestral affinity across populations and geography.** Affinity of contemporary dingoes stratified by population clusters (**A**) and across longitude (**B**). Higher and lower affinity values indicate affinity to the Curracurrang and Nullarbor lineages, respectively.

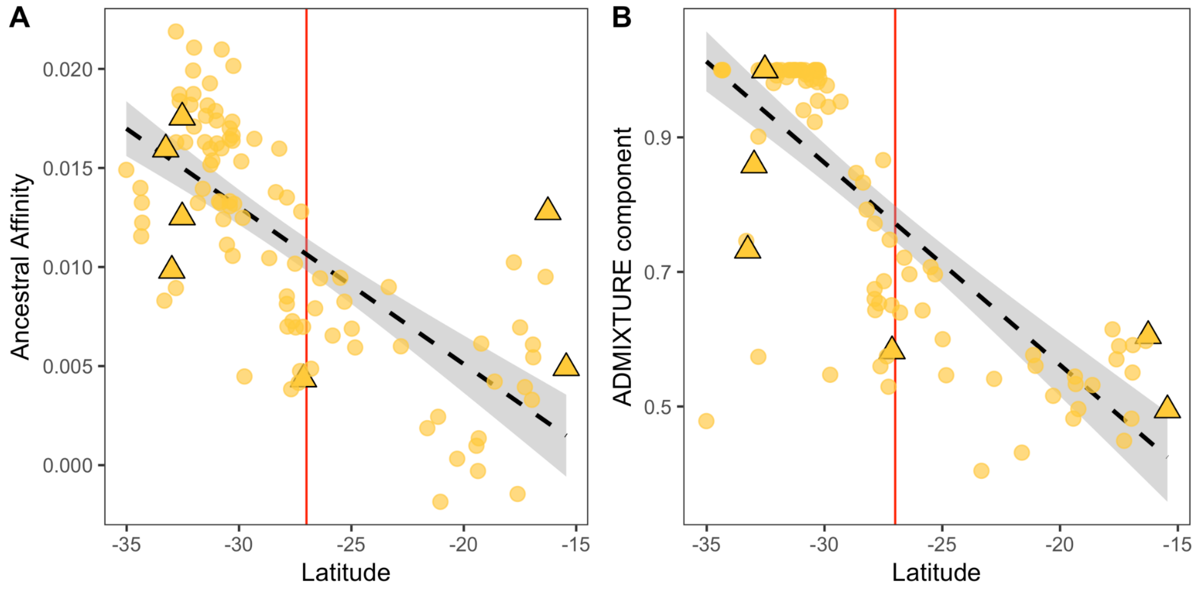


**Figure S17 | Ancestral affinity in the East cluster. A)** Ancestral affinity of dingoes from the East cluster across latitude. The red vertical line indicates the dingo fence, with samples to the left within the fence, and samples to the right north of the fence. **B)** ADMIXTURE component representing the East cluster visualised against latitude.

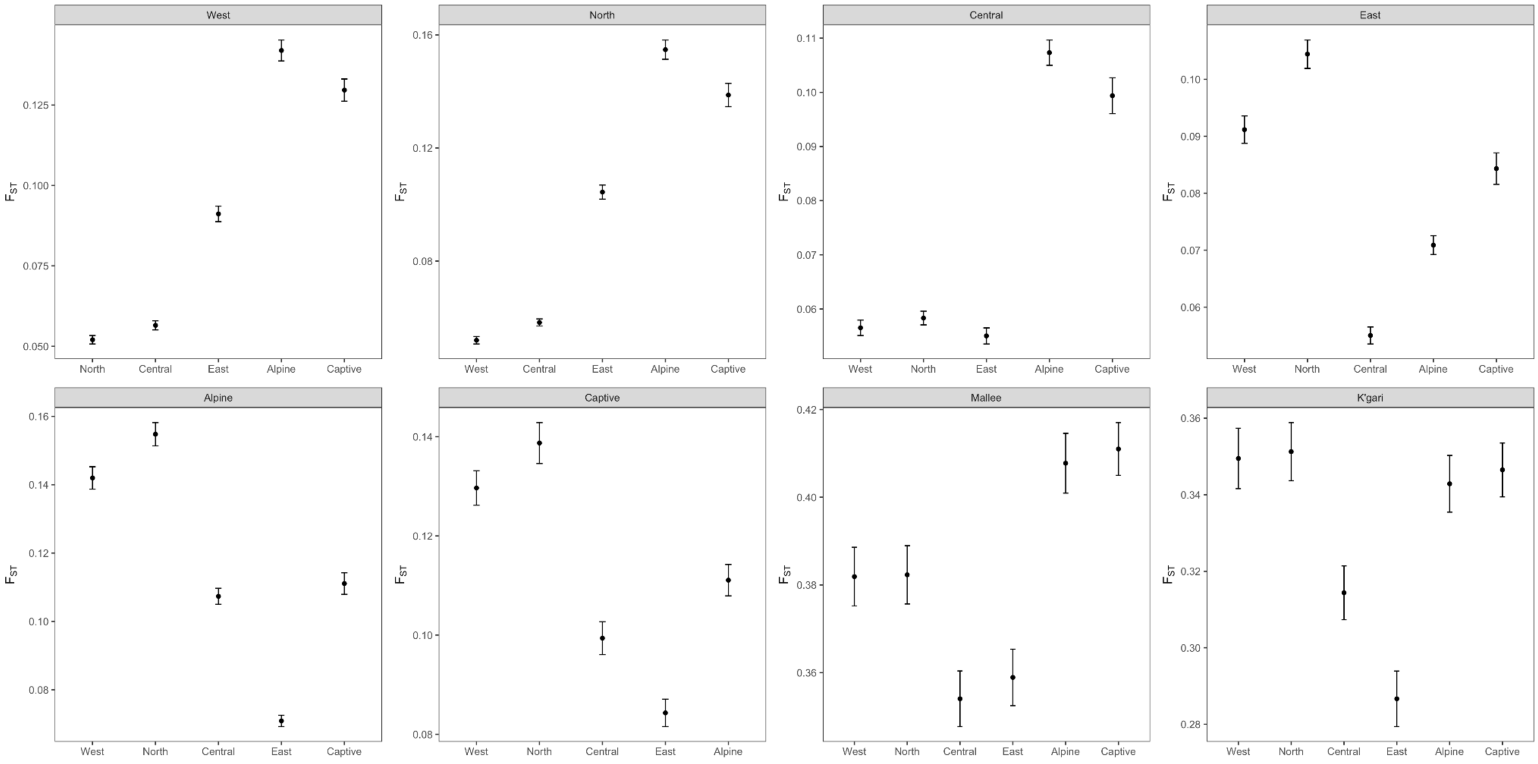


**Figure S18 | $F_{ST}$ between different populations.** Pairwise $F_{ST}$ between dingo populations (facet label vs. x-axis label). The Mallee and K'gari populations were excluded from all panels because their higher genetic differentiation, likely due to small population size, compressed the scale of the plot.

Bergström, A., Frantz, L., Schmidt, R., Ersmark, E., Lebrasseur, O., Girdland-Flink, L., Lin, A.T., Storå, J., Sjögren, K.-G., Anthony, D., Antipina, E., Amiri, S., Bar-Oz, G., Bazaliiskii, V.I., Bulatović, J., Brown, D., Carmagnini, A., Davy, T., Fedorov, S., Fiore, I., Fulton, D., Germonpré, M., Haile, J., Irving-Pease, E.K., Jamieson, A., Janssens, L., Kirillova, I., Horwitz, L.K., Kuzmanovic-Cvetkovic, J., Kuzmin, Y., Losey, R.J., Dizdar, D.L., Mashkour, M., Novak, M., Onar, V., Orton, D., Pasarić, M., Radivojević, M., Rajković, D., Roberts, B., Ryan, H., Sablin, M., Shidlovskiy, F., Stojanović, I., Tagliacozzo, A., Trantalidou, K., Ullén, I., Villaluenga, A., Wapnish, P., Dobney, K., Götherström, A., Linderholm, A., Dalén, L., Pinhasi, R., Larson, G. & Skoglund, P.

(2020). Origins and Genetic Legacy of Prehistoric Dogs. *Science*, 370, 557–564.

Bergström, A., Stanton, D.W.G., Taron, U.H., Frantz, L., Sinding, M.-H.S., Ersmark, E., Pfrengle, S., Cassatt-Johnstone, M., Lebrasseur, O., Girdland-Flink, L., Fernandes, D.M., Ollivier, M., Speidel, L., Gopalakrishnan, S., Westbury, M.V., Ramos-Madrigal, J., Feuerborn, T.R., Reiter, E., Gretzinger, J., Münzel, S.C., Swali, P., Conard, N.J., Carøe, C., Haile, J., Linderholm, A., Androsov, S., Barnes, I., Baumann, C., Benecke, N., Bocherens, H., Brace, S., Carden, R.F., Drucker, D.G., Fedorov, S., Gasparik, M., Germonpré, M., Grigoriev, S., Groves, P., Hertwig, S.T., Ivanova, V.V., Janssens, L., Jennings, R.P., Kasparov, A.K., Kirillova, I.V., Kurmaniyazov, I., Kuzmin, Y.V., Kosintsev, P.A., Lázničková-Galetová, M., Leduc, C., Nikolskiy, P., Nussbaumer, M., O'Drisceoil, C., Orlando, L., Outram, A., Pavlova, E.Y., Perri, A.R., Pilot, M., Pitulko, V.V., Plotnikov, V.V., Protopopov, A.V., Rehazek, A., Sablin, M., Seguin-Orlando, A., Storå, J., Verjux, C., Zaibert, V.F., Zazula, G., Crombé, P., Hansen, A.J., Willerslev, E., Leonard, J.A., Götherström, A., Pinhasi, R., Schuenemann, V.J., Hofreiter, M., Gilbert, M.T.P., Shapiro, B., Larson, G., Krause, J., Dalén, L. & Skoglund, P. (2022). Grey wolf genomic history reveals a dual ancestry of dogs. *Nature*, 607, 313–320.

Botigué, L.R., Song, S., Scheu, A., Gopalan, S., Pendleton, A.L., Oetjens, M., Taravella, A.M., Seregély, T., Zeeb-Lanz, A., Arbogast, R.-M., Bobo, D., Daly, K., Unterländer, M., Burger, J., Kidd, J.M. & Veeramah, K.R. (2017). Ancient European dog genomes reveal continuity since the Early Neolithic. *Nat. Commun.*, 8, 16082.

Bougiouri, K., Aninta, S.G., Charlton, S., Harris, A.C., Petr, M., Carmagnini, A., Piličiauskienė, G., Feuerborn, T.R., Scarsbrook, L., Tabbada, K., Blaževičius, P., Parker, H.G., Gopalakrishnan, S., Larson, G., Ostrander, E.A., Irving-Pease, E.K., Frantz, L.A.F. & Racimo, F. (2025). Imputation of ancient canid genomes reveals inbreeding history over the past 10,000 years. *Proc. Natl. Acad. Sci. U. S. A.*, 122, e2416980122.

Brooks, M.E., Kristensen, K., van Benthem, K.J., Magnusson, A., Berg, C.W., Nielsen, A., Skaug, H.J., Mächler, M. & Bolker, B.M. (2017). glmmTMB Balances Speed and Flexibility Among Packages for Zero-inflated Generalized Linear Mixed Modeling. *R J.*, 9, 378–400.

Browning, B.L., Tian, X., Zhou, Y. & Browning, S.R. (2021). Fast two-stage phasing of large-scale sequence data. *Am. J. Hum. Genet.*, 108, 1880–1890.

Browning, B.L., Zhou, Y. & Browning, S.R. (2018). A one-penny imputed genome from next-generation reference panels. *Am. J. Hum. Genet.*, 103, 338–348.

Cairns, K.M., Crowther, M.S., Parker, H.G., Ostrander, E.A. & Letnic, M. (2023). Genome-wide variant analyses reveal new patterns of admixture and population structure in Australian dingoes. *Mol. Ecol.*, 32, 4133–4150.

Chen, S., Zhou, Y., Chen, Y. & Gu, J. (2018). fastp: an ultra-fast all-in-one FASTQ preprocessor. *Bioinformatics*, 34, i884–i890.

Ferrari, S. & Cribari-Neto, F. (2004). Beta Regression for Modelling Rates and Proportions. *J. Appl. Stat.*, 31, 799–815.

Field, M.A., Yadav, S., Dudchenko, O., Esvaran, M., Rosen, B.D., Skvortsova, K., Edwards, R.J., Keilwagen, J., Cochran, B.J., Manandhar, B., Bustamante, S., Rasmussen, J.A., Melvin, R.G., Chernoff, B., Omer, A., Colaric, Z., Chan, E.K.F., Minoche, A.E., Smith, T.P.L., Gilbert, M.T.P., Bogdanovic, O., Zammit, R.A., Thomas, T., Aiden, E.L. & Ballard, J.W.O. (2022). The Australian dingo is an early offshoot of modern breed dogs. *Sci. Adv.*, 8, eabm5944.

Frantz, L.A.F., Mullin, V.E., Pionnier-Capitan, M., Lebrasseur, O., Ollivier, M., Perri, A., Linderholm, A., Mattiangeli, V., Teasdale, M.D., Dimopoulos, E.A., Tresset, A., Duffraisse, M., McCormick, F., Bartosiewicz, L., Gál, E., Nyerges, É.A., Sablin, M.V., Bréhard, S., Mashkour, M., Bălăşescu, A., Gillet, B., Hughes, S., Chassaing, O., Hitte, C., Vigne, J.-D., Dobney, K., Hänni, C., Bradley, D.G. & Larson, G. (2016). Genomic and archaeological evidence suggest a dual origin of domestic dogs. *Science*, 352, 1228–1231.

Gräler, B., Pebesma, E. & Heuvelink, G. (2016). Spatio-Temporal Interpolation using gstat. *R J.*, 8, 204–218.

Griffith, D.A., Chun, Y. & Li, B. (2019). *Spatial Regression Analysis Using Eigenvector Spatial Filtering*.

Harney, É., Patterson, N., Reich, D. & Wakeley, J. (2021). Assessing the performance of qpAdm: a statistical tool for studying population admixture. *Genetics*, 217, iyaa045.

Li, H. & Durbin, R. (2009). Fast and accurate short read alignment with Burrows–Wheeler transform. *Bioinformatics*, 25, 1754–1760.

Maier, R., Flegontov, P., Flegontova, O., Işıldak, U., Changmai, P. & Reich, D. (2023). On the limits of fitting complex models of population history to f-statistics. *Elife*, 12, e85492.

Manichaikul, A., Mychaleckyj, J.C., Rich, S.S., Daly, K., Sale, M. & Chen, W.-M. (2010). Robust relationship inference in genome-wide association studies. *Bioinformatics*, 26, 2867–2873.

Marcus, J., Ha, W., Barber, R.F. & Novembre, J. (2021). Fast and flexible estimation of effective migration surfaces. *Elife*, 10, e61927.

McInnes, L., Healy, J. & Melville, J. (2018). UMAP: Uniform Manifold Approximation and Projection for Dimension Reduction. *arXiv [stat.ML]*.

McKenna, A., Hanna, M., Banks, E., Sivachenko, A., Cibulskis, K., Kernytsky, A., Garimella, K., Altshuler, D., Gabriel, S., Daly, M. & DePristo, M.A. (2010). The Genome Analysis Toolkit: A MapReduce framework for analyzing next-generation DNA sequencing data. *Genome Res.*, 20, 1297–1303.

Meisner, J., Liu, S., Huang, M. & Albrechtsen, A. (2021). Large-scale inference of population structure in presence of missingness using PCA. *Bioinformatics*, 37, 1868–1875.

Newsome, T.M., Ballard, G.-A., Crowther, M.S., Dellinger, J.A., Fleming, P.J.S., Glen, A.S., Greenville, A.C., Johnson, C.N., Letnic, M., Moseby, K.E., Nimmo, D.G., Nelson, M.P., Read, J.L., Ripple, W.J., Ritchie, E.G., Shores, C.R., Wallach, A.D., Wirsing, A.J. &

Dickman, C.R. (2015). Resolving the value of the dingo in ecological restoration. *Restor. Ecol.*, 23, 201–208.

Plassais, J., Kim, J., Davis, B.W., Karyadi, D.M., Hogan, A.N., Harris, A.C., Decker, B., Parker, H.G. & Ostrander, E.A. (2019). Whole genome sequencing of canids reveals genomic regions under selection and variants influencing morphology. *Nat. Commun.*, 10, 1489.

Rubinacci, S., Ribeiro, D.M., Hofmeister, R.J. & Delaneau, O. (2021). Efficient phasing and imputation of low-coverage sequencing data using large reference panels. *Nat. Genet.*, 53, 120–126.

Salter-Townshend, M. & Myers, S. (2019). Fine-scale inference of ancestry segments without prior knowledge of admixing groups. *Genetics*, 212, 869–889.

Scarsbrook, L., Cairns, K.M., Mitchell, K.J., Bougiouri, K., Evin, A., Harris, A.C., Wood, A.E., Zhang, Z., Lawson, D.J., Alves, J.M., Ditchfield, P.W., Granja-Martins, S., Styring, A.K., Tabbada, K., Thalmann, O., Crowther, M.S., Curry, M., Feuerborn, T.R., Koungoulos, L.G., Letnic, M., Reed, E.H., Sabin, R., Parker, H.G., Ostrander, E.A., Frantz, L.A.F., Larson, G. & Fillios, M.A. (2025). The impacts of European arrival on Australian dingoes. *Proc. Natl. Acad. Sci. U. S. A.*, 122, e2421749122.

Schubert, M., Lindgreen, S. & Orlando, L. (2016). AdapterRemoval v2: rapid adapter trimming, identification, and read merging. *BMC Res. Notes*, 9, 88.

Sinding, M.-H.S., Gopalakrishnan, S., Ramos-Madrigal, J., de Manuel, M., Pitulko, V.V., Kuderna, L., Feuerborn, T.R., Frantz, L.A.F., Vieira, F.G., Niemann, J., Samaniego Castruita, J.A., Carøe, C., Andersen-Ranberg, E.U., Jordan, P.D., Pavlova, E.Y., Nikolskiy, P.A., Kasparov, A.K., Ivanova, V.V., Willerslev, E., Skoglund, P., Fredholm, M., Wennerberg, S.E., Heide-Jørgensen, M.P., Dietz, R., Sonne, C., Meldgaard, M., Dalén, L., Larson, G., Petersen, B., Sicheritz-Pontén, T., Bachmann, L., Wiig, Ø., Marques-Bonet, T., Hansen, A.J. & Gilbert, M.T.P. (2020). Arctic-adapted dogs emerged at the Pleistocene-Holocene transition. *Science*, 368, 1495–1499.

Souilmi, Y., Wasef, S., Williams, M.P., Conroy, G., Bar, I., Bover, P., Dann, J., Heiniger, H., Llamas, B., Ogbourne, S., Archer, M., Ballard, J.W.O., Reed, E., Tobler, R., Koungoulos, L., Walshe, K., Wright, J.L., Balme, J., O'Connor, S., Cooper, A. & Mitchell, K.J. (2024). Ancient genomes reveal over two thousand years of dingo population structure. *Proceedings of the National Academy of Sciences*, 121, e2407584121.

South, A., Michael, S. & Massicotte, P. (2025). *rnaturalearthdata: World Vector Map Data from Natural Earth Used in "rnaturalearth."*

Weeks, A.R., Kriesner, P., Bartonicek, N., van Rooyen, A., Cairns, K.M. & Ahrens, C.W. (2025). Genetic structure and common ancestry expose the dingo-dog hybrid myth. *Evol. Lett.*, 9, 1–12.

Yates, J.A.F., Lamnidis, T.C., Borry, M., Valtueña, A.A., Fagernäs, Z., Clayton, S., Garcia, M.U., Neukamm, J. & Peltzer, A. (2021). Reproducible, portable, and efficient ancient genome reconstruction with nf-core/eager. *PeerJ*, 9, e10947.

Yates, J.A.F., Peltzer, A., Lamnidis, T.C., Borry, M., ZandraFagernas, Bar, I., Valtueña, A.A., alexandregilardet, Bot, N.-C., Garcia, M.U., Floden, E., Ewels, P., Hüther, P., Plessy, C., Gabernet, G., sc13-bioinf, Vågene, Å.J., Patel, H., Botvinnik, O., Carlhoff, S., Hübner, A. & marcel-keller. (2022). *nf-core/eager: [2.4.5] - Wangen (Patch) - 2022-08-02*. Zenodo.

Zhang, S.-J., Wang, G.-D., Ma, P., Zhang, L.-L., Yin, T.-T., Liu, Y.-H., Otecko, N.O., Wang, M., Ma, Y.-P., Wang, L., Mao, B., Savolainen, P. & Zhang, Y.-P. (2020). Genomic regions under selection in the feralization of the dingoes. *Nat. Commun.*, 11, 671.
